# Elab2ARC: A Browser-Based Workspace for Converting Free-Text Protocols into rich FAIR digital objects

**DOI:** 10.64898/2026.05.14.724833

**Authors:** Sabrina Zander, Xiaoran Zhou, Angela Kranz, Kathryn Dumschott, Philippe Rocca-Serra, Heinrich Lukas Weil, Marcel Tschöpe, Timo Mühlhaus, Dirk Von Suchodoletz, Björn Usadel

## Abstract

Electronic laboratory notebooks (ELNs) are widely used in the life sciences, but their notebook format limits machine-readability and FAIR compliance. Consequently, researchers often spend significant manual effort restructuring ELN records into publication-ready outputs. We present elab2ARC, a browser-based workspace that automates the conversion of open-source eLabFTW records into Annotated Research Contexts (ARCs)— version-controlled, ISA-compliant research objects. Using the eLabFTW API, elab2ARC retrieves administrative metadata, protocols, and attachments, reorganising them into ISA-compliant tables and linked datasets. All processing occurs client-side, ensuring user data control before submission to the PLANTdataHUB repository. An optional LLM-assisted workflow extracts structured metadata from free-text protocols, providing editable drafts while preserving human oversight. Designed for use at project completion, elab2ARC reuses existing ELN documentation without disrupting daily laboratory practice. It offers a practical route to FAIR-aligned sharing, publication, and long-term archiving of life-science experimental records.

**Availability and implementation:** elab2ARC is freely accessible at https://nfdi4plants.org/elab2arc/. The source code is available at https://github.com/nfdi4plants/elab2arc under a GPL-3.0 license.

**Supplementary information:** Supplementary data are available online.

## 1. Introduction

Electronic laboratory notebooks (ELNs) are now widely used in the life sciences, replacing paper-based documentation and providing searchable, collaborative storage for experimental protocols, metadata, and associated files (Vandendorpe et al. 2024). In parallel, version control systems such as Git and dedicated data repositories support analysis workflows, change tracking, and collaboration across research teams (Weil et al. 2023).

Despite the broad adoption of these tools, experimental documentation and research data are typically distributed across multiple, loosely connected systems. Protocols and contextual metadata are usually recorded in ELNs as free text. At the same time, large primary datasets (such as sequencing, imaging, or mass spectrometry data) are often stored separately on institutional servers, high-performance computing systems, or cloud infrastructure. When data are prepared for publication, experimental protocols must often be manually rewritten, summarised, and adapted to individual repository requirements. This represents a time-consuming process in which important experimental details are easily lost. As a result, metadata required for interpretation, reuse, and reproducibility are sometimes incomplete or inconsistent in published datasets.

The FAIR principles (Findable, Accessible, Interoperable, Reusable) provide a framework for data management using standardised metadata and machine-readable formats (Wilkinson et al. 2016). In practice, increasing the FAIRness of experimental records remains difficult, because ELNs can be unstructured and are usually not publishable. Linking protocols, metadata, and datasets is therefore a central challenge in research data management. Several metadata standards tackle different metadata layers. Dublin Core (Kunze & Baker 2007) provides a minimal vocabulary for describing digital resources, and Bioschemas (Gray et al. 2017) defines lightweight markup profiles to improve the findability of biological resources. The Investigation–Study–Assay (ISA) framework (Rocca-Serra et al. 2009) provides a domain-independent way to represent experimental workflows, samples, and measurements using tables or JSON. Within ISA, an *Investigation* defines the research context, a *Study* describes the design and samples, and an *Assay* captures specific measurements or analyses. Building on ISA, the Annotated Research Context (ARC) (Weil et al. 2023) uses a metadata-rich RO-Crate packaging format (Soiland-Reyes et al. 2022) that combines ISA-based metadata, data files, and documentation into a single version-controlled research object, thereby enabled provenance tracking and reproducible data publication. Completed ARCs can subsequently be published in FAIR public repositories, e.g. the PLANTdataHUB (Weil et al. 2023), bridging local data management and public access.

To bridge ELNs and FAIR data management, several tools aim to improve interoperability between ELNs and structured metadata frameworks. While some ELNs, including eLabFTW, can export complete entries as .eln files, a format based on the RO-Crate specification (Soiland-Reyes et al. 2022), these exports preserve the original document structure rather than restructuring it into experimental metadata. LISTER (Musyaffa et al. 2023) extracts ISA-compliant metadata from manually annotated eLabFTW entries, and Schröder et al. (Schröder et al. 2022) proposed a structure-based method that translates ELN layouts into semantic representations for inclusion in RO-Crates. However, to the best of our knowledge, no lightweight solution currently converts ELN entries into structured, version-controlled, FAIR-compliant research objects while preserving detailed protocols and their links to the associated datasets. Bridging this gap typically falls to the researchers responsible for preparing data for publication.

Here, we present elab2ARC, a browser-based workspace that converts experimental records from the free and open-source ELN eLabFTW into ARCs. eLabFTW is widely used in life science laboratories as a flexible, open-source ELN that accommodates heterogeneous documentation while not enforcing a strict internal data structure (CARPi et al. 2017). The tool retrieves metadata, protocols, and attachments through the eLabFTW API and assembles them into ISA-compliant, Git-versioned ARCs entirely in the user’s browser. Optionally, it applies either local or API-accessed large language models (LLMs) to extract structured metadata from free-text protocols, reducing manual annotation effort. By integrating ELN documentation, standardised metadata, and version-controlled repositories in a single workflow, elab2ARC supports reproducible and FAIR-aligned data management without requiring researchers to change their day-to-day documentation habits.

## 2. Methods

### System overview

Elab2ARC supports ELN users who want to apply FAIR Digital Objects for data management. It performs a one-way conversion of eLabFTW records into an existing or new ARC, mapping administrative metadata and protocols to ISA-compliant tables and Markdown (González-Beltrán et al. 2014). The tool focuses on generating standards-based objects for further processing or release via PLANTdataHUB (Weil et al. 2023). A core design principle is its browser-based operation; all data retrieval, transformation, and Git operations occur locally or via authenticated API calls. This ensures data to be under user’s control without requiring additional server-side infrastructure or local installations.

### Software architecture

The interface of elab2ARC highlights user interaction and hides automated processing (Figure 1). Users authenticate via tokens, select eLabFTW records, and specify a target ARC. Subsequent steps are automated: retrieving metadata and files via API, managing them in memfs, and assembling the ISA-compliant hierarchy. Finally, the ARC is committed to an in-memory Git repository and pushed to PLANTdataHUB (Weil et al. 2023).

**Figure 1.**
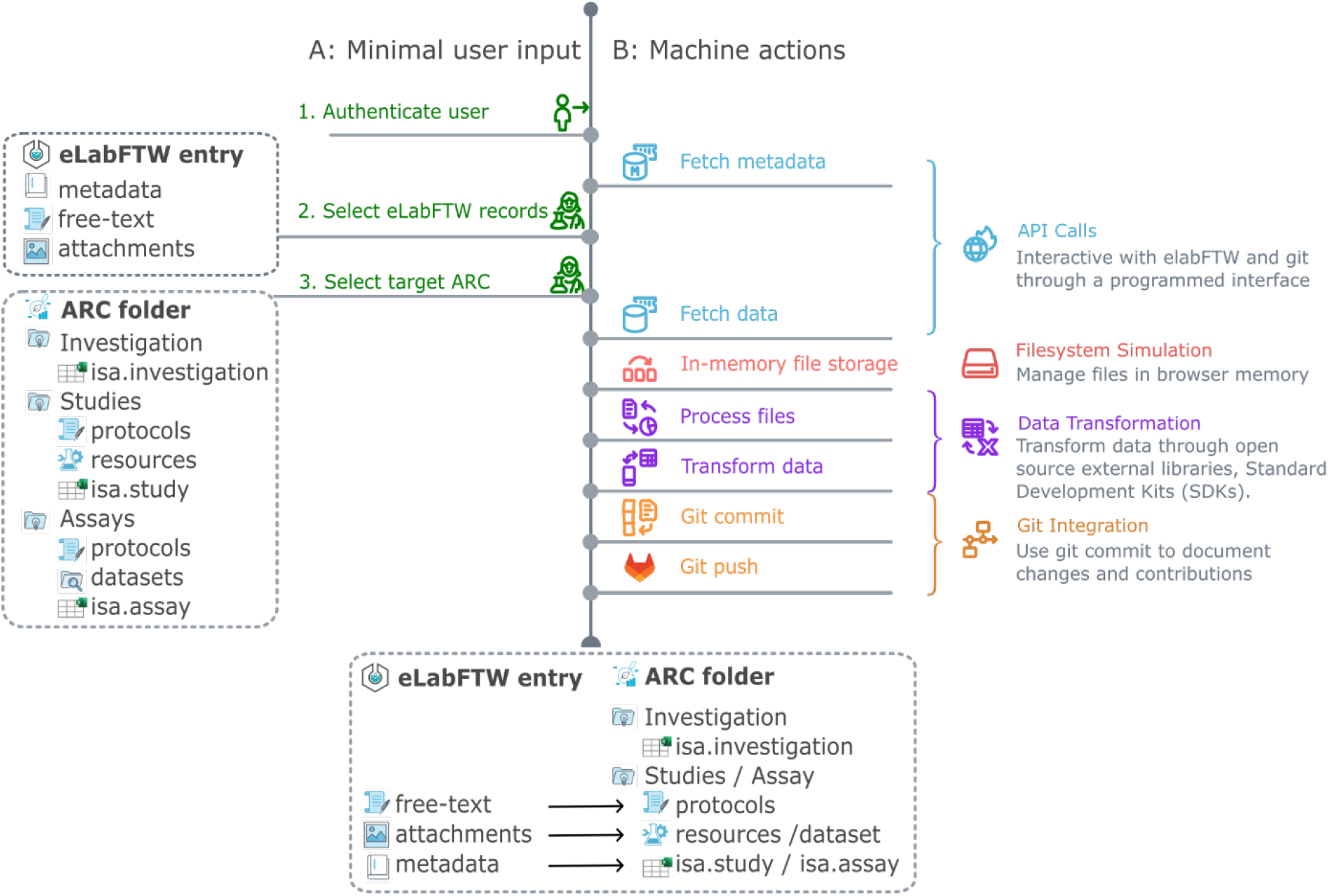
User input and automated processing steps in the elab2ARC workflow. The figure illustrates the elab2ARC workflow, distinguishing between minimal user input and automated machine actions. Users perform only three actions: authentication, selecting eLabFTW records, and selecting a target ARC. All subsequent steps— including data retrieval, in-memory file handling, metadata transformation into ISA/ARC-compliant structures, and Git-based version control—are executed automatically on the client side. The ARC block represents parts of the standard ARC directory structure generated during the conversion process.

### Integration with eLabFTW

Data of experiments and resources can both be retrieved by elab2ARC through eLabFTW API. The data is sorted into three categories (i) top-level administrative metadata, (ii) experiment-notes as free-text protocols, and (iii) attached files such as images or spreadsheets (Figure S1). Both experiments and resources are supported because eLabFTW uses resources to represent reusable items, such as samples, reagents, and equipment, that are commonly referenced in experiments.

### Mapping of eLabFTW metadata to the ISA framework

eLabFTW entries separate administrative metadata—like identifiers, authorship, and tags— from free-text content. The elab2ARC tool transforms this data into a structured representation based on the ISA framework. Selected entries are mapped to ARC components: metadata is written to ISA-compliant isa.study or isa.assay files, while protocols are converted to Markdown and stored in protocol directories. This preserves the logical association between metadata, protocols, and datasets. The transformation uses ARCtrl, a library for manipulating ARCs and ISA-XLSX files, ensuring alignment with community standards and supporting long-term interoperability.

### Version control and repository integration

Generated ARCs are managed as Git repositories. During conversion, all generated files (ISA metadata tables, protocol documents, and attachments) are staged and committed to the in-memory Git repository. Each conversion produces a reproducible snapshot of the ARC derived from the selected entries. Files located in any dataset folder are handled automatically through Git Large File Storage. The repository is then pushed to PLANTdataHUB. The public ARC can later be released as a FAIR publication with an assigned Digital Object Identifier, as described in (Weil et al. 2023).

### Optional LLM-assisted protocol structuring

ELNs contain free-text because it is a notebook by design, which prioritizes convenience overformality. To increase interoperability, elab2ARC provides an optional LLM-assisted workflow that extracts structured metadata from free-text ELN notes. The LLM identifies entities like samples, inputs, and parameters, writing them to ISA-compliant annotation tables in isa.study or isa.assay files. The LLM model and prompt can both be customized via web interface (Supplementary Files S1-S3). Importantly, the process is transparent: generated metadata serves as a refinement starting point, ensuring researchers maintain final control over annotations before they are integrated alongside manually extracted metadata.

### Implementation and availability

elab2ARC is a client-side web application built with JavaScript, HTML, and CSS that runs entirely in modern browsers without a backend. It utilizes ARCtrl for ISA-XLSX handling, isomorphic-git for Git operations, and memfs for in-memory file storage. The tool interacts with external systems, like eLabFTW and the PlantdataHUB, solely through authenticated API calls using personal access tokens. Released as open-source software, elab2ARC is hosted at nfdi4plants.org/elab2arc/. Comprehensive documentation and the source code are available via GitHub and the DataPLANT Knowledge Base, ensuring transparency and ease of use for the community.

## 3. Results

### 3.1 Use case: Conversion of a genomic workflow from eLabFTW into ARC

We demonstrate elab2ARC using a genomic workflow consisting of five eLabFTW experiments: Bacterial cultivation, DNA Extraction, Library Preparation, Sequencing Run, and Bioinformatic Analysis. Documented as free-text protocols with administrative metadata and attachments, these records are available on a demo instance (https://elab.dataplan.top/) and as Supplementary Files S4-S8. The use case illustrates how elab2ARC consolidates these distributed materials—including references to external raw data—into a structured, reusable research object suitable for collaboration or formal publication (Figure 2).

**Figure 2.**
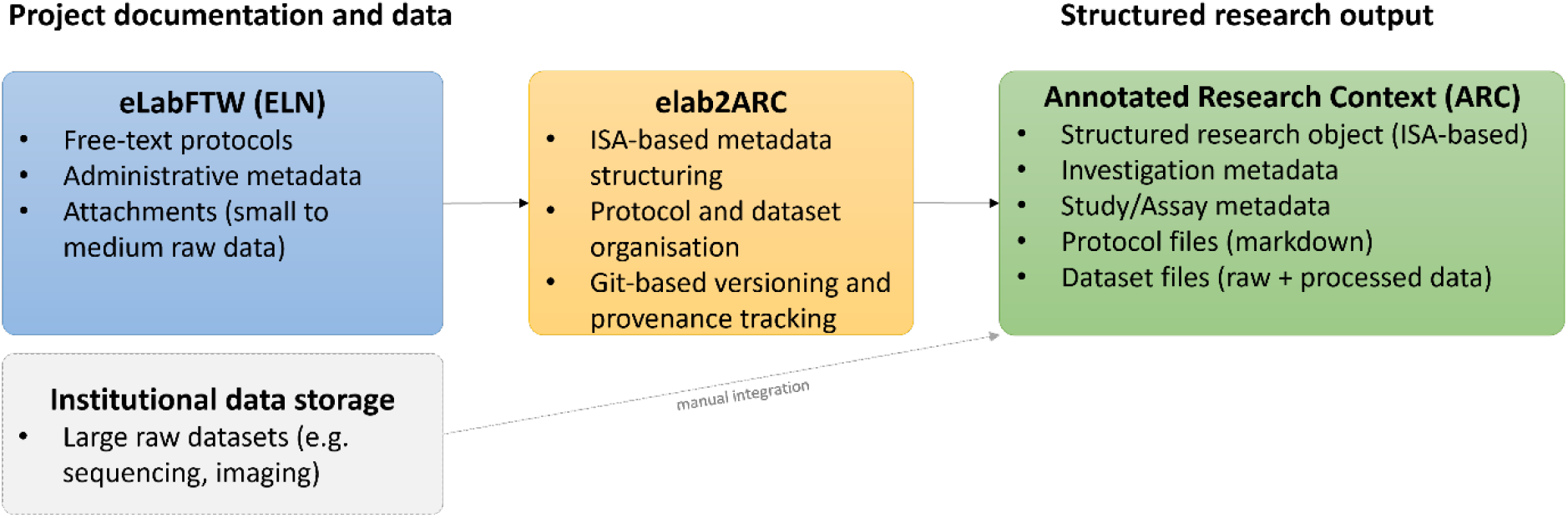
Overview of the elab2ARC use case. Experimental records documented in eLabFTW are automatically converted by elab2ARC into a structured ARC. Free-text protocols, administrative metadata, and attached data files are reorganised into ISA-compliant metadata tables, protocol files, and datasets under Git-based version control. Large raw datasets stored on institutional infrastructure are manually consolidated into the ARC. The resulting ARC provides a single, structured research object for reuse, sharing, and publication.

Elab2ARC (https://nfdi4plants.org/elab2arc/) requires an access token to establish a connection to the source eLabFTW instance and one for the PLANTdataHUB. On the token page, the researcher first selects their eLabFTW instance. For this use case, the demo instance (https://elab.dataplan.top/) is selected (Figure 3, step 1). Clicking “Get a token” automatically retrieves and fills the corresponding API token without requiring manual input (Figure 3, step 2). Researchers connecting their own institutional eLabFTW instance can alternatively provide a self-generated API token, which can be created in the eLabFTW user settings. A token is required to create or modify ARCs on the PLANTdataHUB (Figure 3, step 3). If the researcher is already logged into the PLANTdataHUB in their browser, clicking “Get a token” retrieves the token automatically. For the demo, a shared DataHUB account is available using the credentials provided in the Data Availability section. After saving the tokens (Figure 3, step 4), the researcher selects the target experiments from a list of all available ELN entries. In this use case, five experiments were selected (Figure 3, step 5a/b) and confirmed (Figure 3, step 6). In the last step, a new ARC is created (Figure 3, step 7), or a target ARC and the target directory within the ARC, in our use case the assay folder, is selected (Figure 3, step 7b). Finally, the conversion process is started (Figure 3, step 8). These three user interactions constitute the only required manual steps in the workflow; all subsequent processing steps are performed automatically (Figure 1).

**Figure 3.**
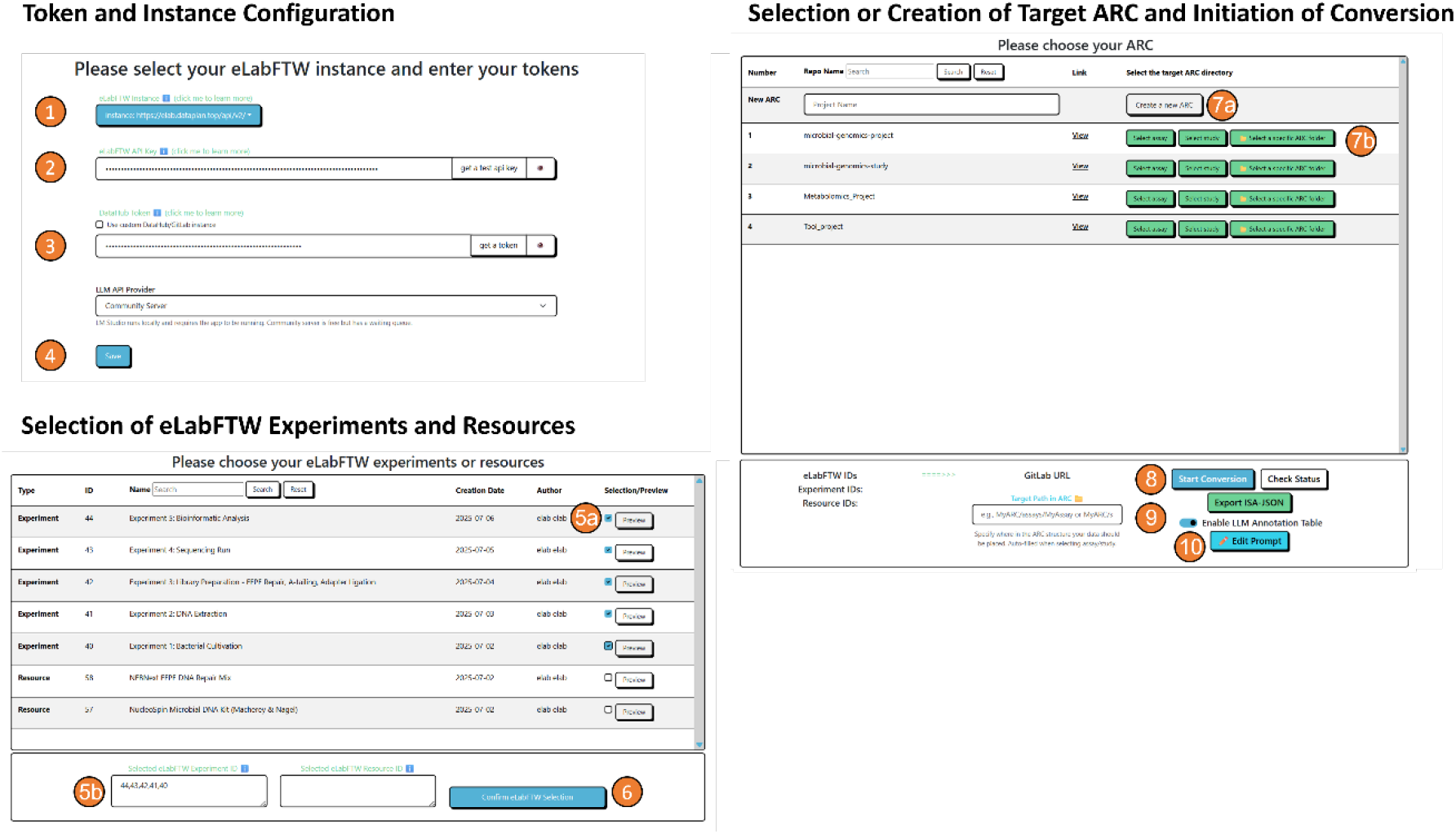
User interaction steps in the elab2ARC web interface. Screenshots illustrate the three required user actions for converting eLabFTW records into an ARC. (1–4) Token and instance configuration by selecting an eLabFTW instance, providing API tokens for eLabFTW and PLANTdataHUB, and saving the configuration. (5a–6) Selection of eLabFTW experiments and resources via the integrated table view, followed by confirmation of the selection. (7a–10) Selection or creation of a target ARC, specification of the target directory within the ARC, optional activation of LLM-assisted annotation and prompt editing, and initiation of the conversion process. Only interface components relevant for the required user actions are shown.

Upon user input, elab2ARC automatically retrieves selected ELN records and converts them into a structured ARC. Each experiment maps to a dedicated assay, where the title determines the folder name to ensure traceability. Using Experiment 2 - DNA Extraction as an example (Figure S1), the process converts HTML protocols into Markdown for the protocols/ directory, while attachments like gel images move to the dataset/ directory. Top-level metadata (including authorship, timestamps, and identifiers) are recorded in the isa.assay.xlsx file. The completed ARC is pushed to the PLANTdataHUB, creating a machine-actionable snapshot of the experimental context. This repository (accessible at git.nfdi4plants.org/elab/Genomics) serves as a foundation that can be enriched with raw sequencing data and further metadata, or used directly for data publication and long-term archival.

### 3.2 Optional protocol structuring using LLM assistance

When elab2ARC is run without LLM support, the generated isa.assay.xlsx adheres to the ISA-Tab standard and includes the top-level metadata (Figure S2). When optional LLM-assisted extraction is enabled, elab2ARC sends the free-text protocol to a locally or externally hosted LLM, which can be configured on the token page after enabling the LLM option (Figure 3, step 9). Using a predefined prompt (Supplementary File S3), which is editable by the user (Figure 3, step 10), the LLM identifies experimental parameters, sample characteristics, and process steps from the unstructured text and returns structured data that is written to additional sheets within the same isa.assay.xlsx file. For Experiment 1 (Bacterial cultivation), the LLM successfully extracted information about the samples and experimental parameters (Supplementary Files S9-S10). An example of an LLM-generated ISA assay table derived from a free-text protocol is shown in Supplementary Figure S2.

The LLM-assisted workflow reduces the manual effort of structuring free-text protocols while maintaining transparency. Researchers retain control, as all generated metadata can be reviewed or edited. However, extraction quality depends on the original ELN documentation. In Experiment 3 (Library Preparation), unstructured text led to poor parameter extraction, highlighting that elab2ARC is a conversion tool, not a substitute for good documentation practices. It only extracts what is explicitly recorded.

## Conclusion

ELNs are widely adopted as primary systems for documenting experimental work in the life sciences (Vandendorpe et al. 2024). However, their document-oriented design is optimised for human-readable record keeping rather than for structured research data management or machine-actionable reuse. As a result, substantial manual effort is often required to consolidate experimental records, extract relevant metadata, and prepare data for sharing or publication. The results presented here demonstrate how elab2ARC addresses this gap by providing a bridge between ELN-based documentation and standards-based research objects. elab2ARC enables semi-automated transformation of ELN entries into version-controlled FAIR research objects without changing lab practices. By mapping experiments to ISA-aligned ARCs, it preserves context while ensuring machine-actionability (González-Beltrán et al. 2014; Rocca-Serra et al. 2009). This approach establishes explicit links between data and protocols, significantly reducing manual publication effort. elab2ARC consolidates and curates ELN records for reuse without enforcing rigid workflows. This approach maintains usability in diverse labs, assisting researchers only when preparing data for dissemination and publication (Higgins et al. 2022). Unlike annotation-driven tools like LISTER that require predefined tags and strict syntax during documentation, elab2ARC reduces researcher burden by transforming existing eLabFTW content into ISA-compliant representations without mandatory prior structuring (Musyaffa et al. 2023). Solutions such as ELNdataBridge (Starman et al. 2025) connect multiple ELN systems via server-centric middleware and predefined mappings. While these approaches facilitate cross-institutional data exchange, they rely on centralised services and fixed pipeline integrations, which can reduce flexibility and increase infrastructural complexity in heterogeneous research environments. More general metadata packaging approaches, including RO-Crate (Soiland-Reyes et al. 2022), provide lightweight, machine-readable containers for research data and their metadata. ELNs such as eLabFTW already support export into RO-Crate–compatible formats, enabling standardised packaging of experimental records. However, these exports largely preserve the original document structure and do not address the transformation of free-text protocols into structured, ISA-based experimental metadata. Consequently, additional curation steps are still required to achieve fully machine-actionable representations suitable for downstream analysis and repository submission. In contrast, elab2ARC builds directly on existing ELN content and performs a semantic reorganisation into ISA-aligned ARC components without requiring prior structuring or annotation. This reduces duplication of effort and lowers the entry barrier for FAIR-aligned research data management.

The optional LLM-assisted workflow demonstrates how AI can complement research data management standards. By extracting metadata from narrative content, it reduces the manual effort required to create ISA-compliant records. Following recent studies, elab2ARC treats LLM-generated output as a transparent starting point rather than authoritative data. This maintains human oversight and allows for manual refinement, mitigating risks associated with automated curation while ensuring high-quality, machine-actionable annotations (Tan et al. 2024).

While elab2ARC effectively converts eLabFTW records into structured ARCs, some limitations remain. The quality and completeness of generated metadata depend on the consistency and detail of ELN documentation, an issue widely reported in ELN-based workflows (Higgins et al. 2022). Highly heterogeneous or sparse protocol descriptions may still require manual refinement after conversion.

## Acknowledgements

We would like to thank our colleagues from MibiNet, CEPLAS, SNP2Prot, and DataPLANT for their valuable feedback and suggestions to improve the elab2ARC tool. In particular, we would like to acknowledge Jenny Köpplin, Vanessa Uglosch, Dr. Dominik Brilhaus, and Dr. Dennis Psaroudakis for discussion and testing.

## Supplementary data

Supplementary data are available online.

## Data availability

The elab2ARC tool is freely accessible via the DataPLANT web interface at https://nfdi4plants.org/elab2arc/. The underlying source code is available in the public GitHub repository at https://github.com/nfdi4plants/elab2arc under an open-source license. Documentation and user guidance are provided through the DataPLANT Knowledge Base at https://nfdi4plants.github.io/nfdi4plants.knowledgebase/resources/elab2arc.

One-click access to elab2arc, as well as usernames and passwords to access our demo eLabFTW and DataHUB, can be found at https://elab2arc-review.dataplan.top. This example ARC is also available here: https://git.nfdi4plants.org/elab/Genomics.

## References

CARPi, N., Minges, A. & Piel, M.,2017. eLabFTW: An open source laboratory notebook for research labs. Journal of Open Source Software, 2(12), p.146.

González-Beltrán, A. et al.,2014. linkedISA: semantic representation of ISA-Tab experimental metadata. BMC Bioinformatics, 15 Suppl 14(S14), p.S4.

Gray, A., Goble, C. & Jiménez, R.,2017. Bioschemas: From potato salad to protein annotation.International Workshop on the Semantic Web.

Higgins, S.G., Nogiwa-Valdez, A.A. & Stevens, M.M.,2022. Considerations for implementing electronic laboratory notebooks in an academic research environment. Nature Protocols, 17(2), pp.179–189.

Kunze, J. & Baker, T.,2007. The Dublin Core Metadata Element Set, RFC Editor. Available at: 10.17487/rfc5013.

Musyaffa, F.A., Rapp, K. & Gohlke, H.,2023. LISTER: Semiautomatic metadata extraction from annotated experiment documentation in eLabFTW. Journal of Chemical Information and Modeling, 63(20), pp.6224–6238.

Rocca-Serra, P.Sansone, S.-A. & Brandizi, M.,2009. Specification documentation: ISA-TAB 1.0, Available at: 10.5281/zenodo.161355.

Schröder, M. et al., 2022. Structure-based knowledge acquisition from electronic lab notebooks for research data provenance documentation. Journal of Biomedical Semantics, 13(1), p.4.

Soiland-Reyes, S. et al., 2022. Packaging research artefacts with RO-Crate. Data Science, 5(2), pp.97–138.

Starman, M. et al., 2025. ELNdataBridge: facilitating data exchange and collaboration by linking Electronic Lab Notebooks via API. Journal of Cheminformatics, 17(1), p.86.

Tan, Z. et al., 2024. Large Language Models for data annotation and synthesis: A survey. arXiv [cs.CL]. Available at: 10.48550/arXiv.2402.13446.

Vandendorpe, J. et al., 2024. Ten simple rules for implementing electronic lab notebooks (ELNs). PLoS computational biology, 20(6), p.e1012170.

Weil, H.L. et al., 2023. PLANTdataHUB : a collaborative platform for continuous FAIR data sharing in plant research. The Plant Journal, 116(4), pp.974–988.

Wilkinson, M.D. et al., 2016. The FAIR Guiding Principles for scientific data management and stewardship. Scientific data, 3(1), p.160018.

