## Supplementary figures and images for "Elab2ARC: A Browser-Based Workspace for Converting Free-Text Protocols into rich FAIR digital objects"

### FigureS1

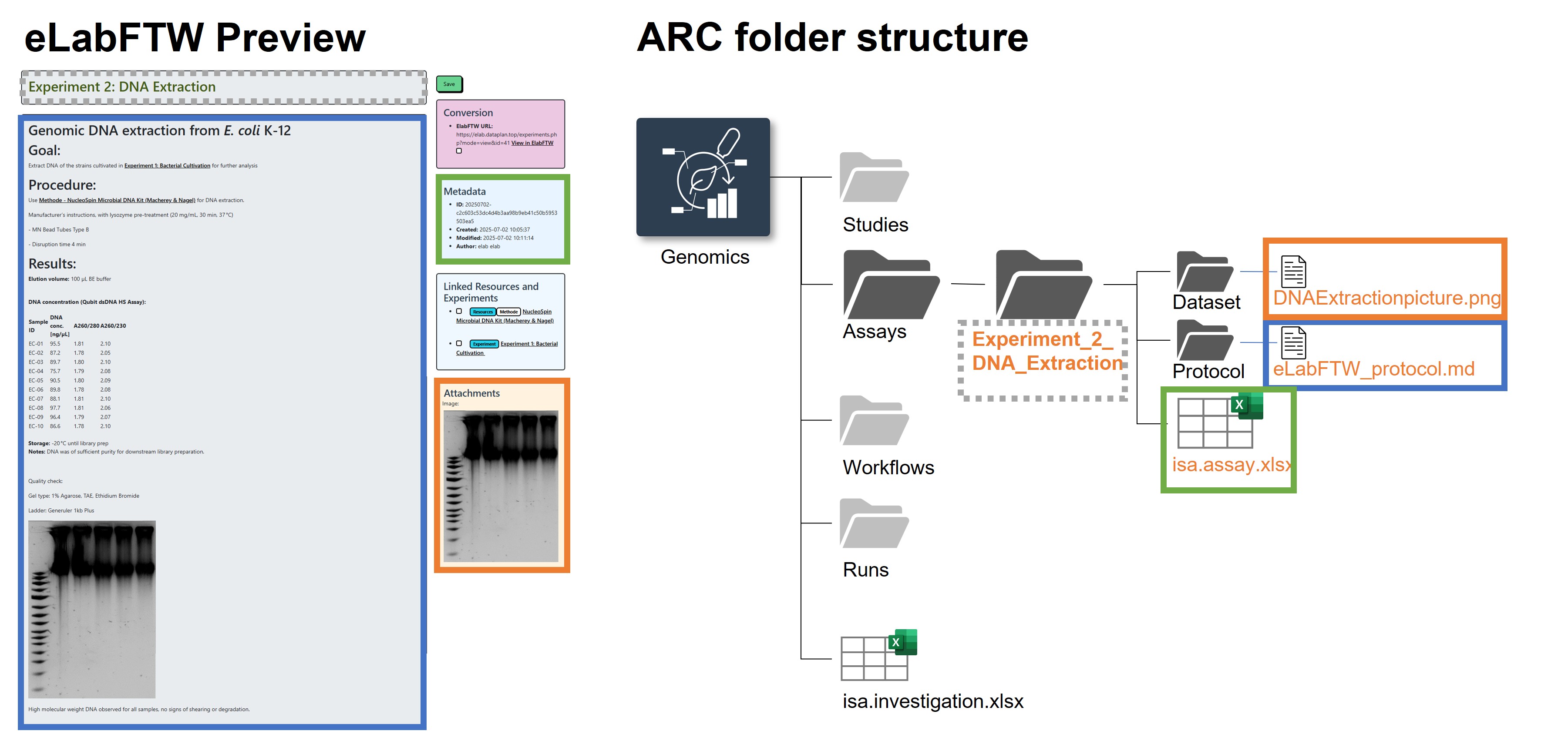

### FigureS2

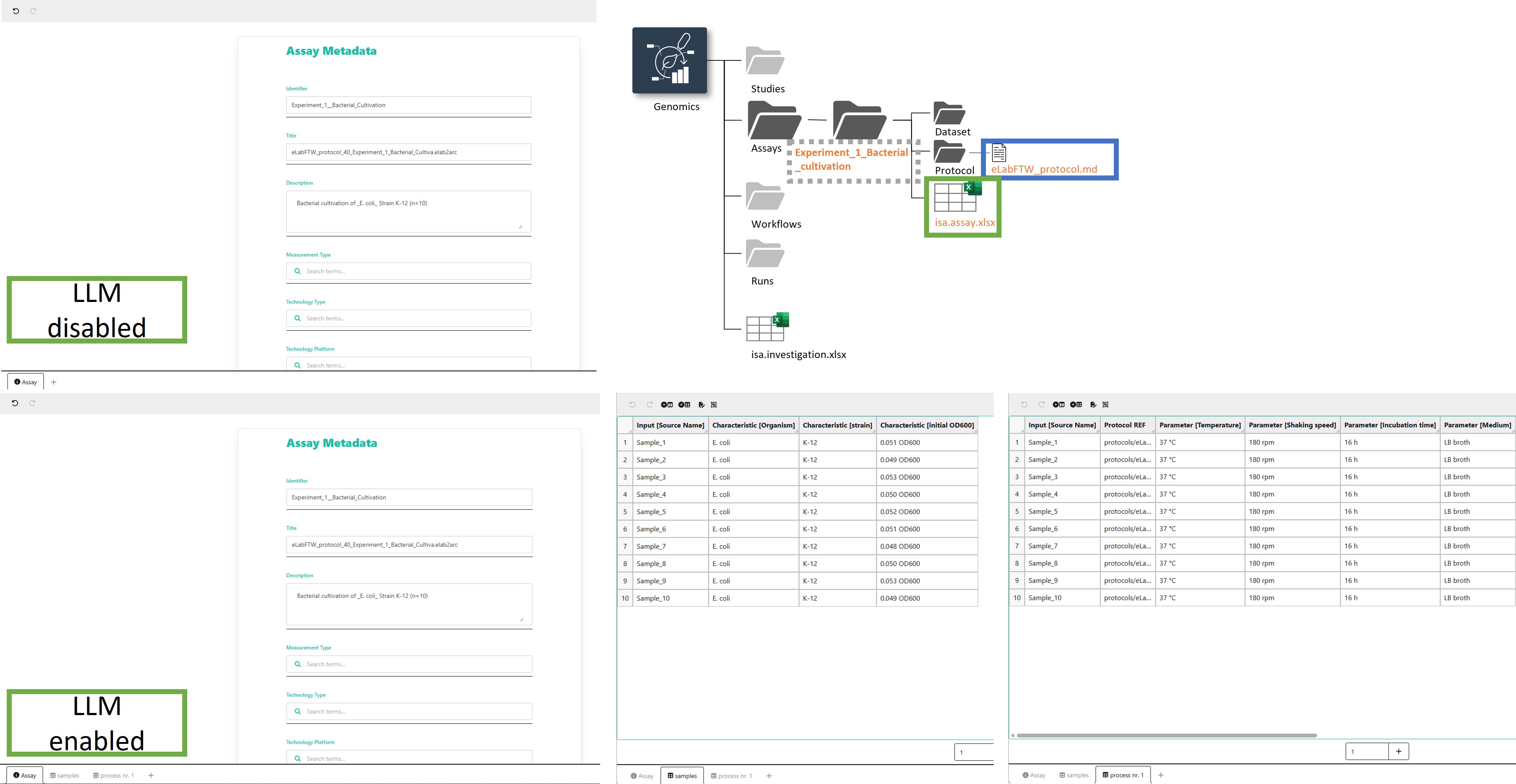
