## Supplementary material for "Elab2ARC: A Browser-Based Workspace for Converting Free-Text Protocols into rich FAIR digital objects": S1_Supplementary_File

### Supplementary Material: LLM Function Design and Configuration

*Protocol-to-ISA Conversion: Large Language Model Integration*

#### 1. Introduction

This document describes the large language model (LLM) integration within the elab2ARC conversion pipeline, which transforms unstructured experimental protocols from eLabFTW into structured ISA-Tab (Investigation, Study, Assay) and ARCs (annotated research contexts). The LLM function is responsible for semantic extraction of samples, protocols, parameters, and data file references from free-text markdown protocols. The design prioritizes reproducibility, provider structured and deterministic output, and graceful degradation when AI services are unavailable. All configuration details, prompt engineering decisions, and resilience mechanisms are documented here to ensure full transparency for peer review and replication.

#### 2. API Provider Architecture

The LLM service module implements a multi-provider architecture that abstracts API differences behind a unified interface. This design decision ensures that researchers are not locked into a single commercial provider and can operate entirely within institutional infrastructure or offline environments. Five provider profiles are pre-configured:

- **DataPlan (Default) —** Institution-hosted proxy requiring no API key; adds Host and institution headers for routing. This is the default because it removes friction for new users and ensures no credentials are transmitted to third-party services.
- **Together.AI —** Commercial inference platform with access to high-performance open-weight models. Requires a Bearer token. Selected when users need models not available locally or when the institutional proxy is unreachable.
- **LM Studio (Local) —** Desktop application for local model serving. Ideal for air-gapped environments or protocols containing sensitive data that cannot leave the workstation. No API key required.
- **Ollama (Local) —** Lightweight command-line tool for running models locally. Complements LM Studio with a simpler setup path favoured by power users and CI/CD pipelines.
- **Custom Endpoint —** Arbitrary OpenAI-compatible API endpoint. Supports internal Kubernetes deployments, vLLM stacks, or emerging providers without waiting for a client-side update.

Provider selection is persisted in browser localStorage, so returning users retain their preferred backend without reconfiguration. The modular header builder injects provider-specific authentication (Bearer tokens for Together.AI, Host and institution tags for

DataPlan) while always enforcing *Content-Type: application/json*. This abstraction allows the same prompt and parsing logic to run regardless of whether the model executes on a remote GPU cluster or a local Apple-Silicon laptop.

#### 3. Model Selection and Configuration

##### 3.1 Supported Models

The system maintains a curated allow-list of models (VALID_MODELS) to prevent users from selecting incompatible or deprecated endpoints. Two production-grade models are currently supported:

- **Qwen/Qwen3-235B-A22B-Instruct-2507-tput (Default).** 131k-token context window; Mixture-of-Experts (MoE) architecture with 235 billion total parameters and 22 billion active parameters per forward pass. Selected as default after benchmarking 350 protocols because it achieved the highest success rate (100 %) and zero hallucination on the dataFiles field after prompt hardening.
- **openai/gpt-oss-120b.** 32,768-token context window; 120-billion-parameter dense transformer. Serves as a fallback when the primary model is rate-limited or unavailable. Its smaller context window makes it more suitable for short protocols or when latency is critical.

Model identifiers are validated against the allow-list on every load; if an invalid identifier is found in localStorage (for example, after a model is retired), it is automatically cleared and the default is restored. This prevents cryptic API errors from stale configuration.

##### 3.2 Context Window Management

Each model advertises a fixed context window (131K for Qwen3-235B, 32K for GPT-OSS-120B). The service reserves 1,000 tokens for the prompt template itself (system instructions, schema definition, and examples) and an additional 2,000 tokens for the model response. The remaining capacity is the maximum input chunk size. If a protocol exceeds this budget, hierarchical chunking is triggered automatically. This conservative reservation avoids mid-generation truncation, which is a common cause of unparseable partial JSON in streaming mode.

#### 4. Prompt Engineering Design

The extraction prompt is assembled from four independently editable sections stored in browser localStorage. This modular architecture lets advanced users tune specific aspects of the prompt without risking accidental corruption of the JSON schema or examples. When no custom prompt exists, a hardened default is used.

- **System Role:** Defines the persona ("scientific data extraction assistant") and the high-level task: analyse a protocol and emit ONLY a JSON object with no markdown, no explanation, and no conversational filler. Constraining the model to a single machine-readable output format eliminates post-processing regex that would otherwise be fragile across model versions.

- **JSON Schema:** Specifies the exact output structure with nested arrays for samples, protocols, parameters, and dataFiles. Each field is annotated with inline type hints (e.g., "array of input sample/material names — ONE VALUE PER ROW"). These hints act as soft constraints that reduce positional errors without consuming the formal token budget of a JSON Schema draft.
- **Extraction Rules:** Contains domain-specific imperatives: extract ALL parameters including software versions and command-line flags; link protocols sequentially so that the output of step N exactly matches the input of step N+1; enforce array-length parity across inputs, outputs, and dataFiles. The rules section is the primary lever for hallucination mitigation (see Section 6.3).
- **Examples:** Provides five concrete, annotated examples covering good parameter extraction, protocol chaining, sample characteristics with ontology term sources, and data file duplication patterns (shared files vs. per-sample files). In-context learning through examples outperforms zero-shot extraction for specialised scientific vocabulary.

A full-screen Prompt Editor modal exposes each section in a dedicated tab, with a read-only preview tab showing the assembled prompt exactly as it will be transmitted. Version history stores the last 50 saved configurations, enabling diff-based comparison and one-click rollback. This audit trail is essential for reproducible research: if a conversion result changes, the metadata record (Section 9) captures which prompt version was active.

#### 5. API Parameters and Their Rationale

Every LLM request is dispatched with a fixed parameter set chosen to maximise structured-output fidelity while minimising latency and cost.

- **temperature = 0.1.** A near-greedy sampling temperature suppresses creative paraphrasing and enforces lexical consistency. In structured extraction tasks, high temperatures increase the risk of field omissions, synonym substitutions (e.g., "incubation time" vs. "incubation duration"), and spurious markdown wrappers. The value 0.1 strikes a balance: it is low enough to make repeated extractions of the same protocol nearly identical, yet high enough to allow minor rephrasing when the source text is ambiguous.
- **max_tokens = 8,192.** The output ceiling is set well above the typical response size (~1,500 tokens for a three-step protocol) to accommodate long protocols with many parameters and samples. Early prototypes used 4,096 tokens, which caused truncation in roughly 8 % of bioinformatics pipelines that enumerate dozens of software parameters. Doubling the limit eliminated truncation without materially increasing cost, because the model stops at the end of the JSON object.
- **stream = true.** Streaming is enabled for real-time user feedback. Token chunks are appended to a debug accordion panel as they arrive, giving users immediate visibility

into model reasoning. Streaming also improves perceived latency: the first tokens appear within seconds even if the full response requires 30–60 seconds.

- **enable_thinking = false (local providers only).** LM Studio and Ollama models occasionally emit chain-of-thought reasoning text before the JSON payload. This preamble breaks deterministic regex extraction. The service explicitly disables thinking mode for local providers, forcing the model to emit the JSON object from the first token. Remote providers (DataPlan, Together.AI) handle this server-side.

Notably, the system does not expose top_p or frequency_penalty to end users. In extraction tasks, these parameters correlate with output instability rather than quality. Removing them from the UI reduces the configuration space and prevents accidental degradation of results.

#### 6. Text Chunking Strategy

Long experimental protocols can exceed the available context window after prompt overhead is reserved. Rather than silently truncating text, which guarantees information loss, the service implements a hierarchical chunking algorithm that respects natural language boundaries.

- **Section-level split:** The protocol is first divided at Markdown heading boundaries (### and ##). Headings demarcate semantic units (e.g., "Sample Preparation", "DNA Extraction", "Sequencing"), so keeping each unit intact preserves local context. Sections are greedily packed into chunks without exceeding the token budget.
- **Paragraph-level split:** If an individual section is larger than the chunk limit, it is subdivided at blank-line boundaries (\n\n+). Paragraphs typically describe a single operation or condition, making them the next-best atomic unit.
- **Sentence-level split:** If a single paragraph still exceeds the limit, sentence boundaries (. ! ?) are used. This is a last resort because it may sever causal clauses (e.g., "Incubate at 37 °C for" / "30 minutes"), but it guarantees that no chunk is dropped.
- **Forced character split:** In the degenerate case of a sentence longer than the budget, the text is hard-split at the character limit. This occurs rarely (<0.1 % of protocols) and is logged as a warning.

Token estimation uses a conservative heuristic of one token per four characters. While exact tokenisation requires model-specific vocabulary tables, the 4:1 ratio is empirically safe for English scientific text and introduces a small buffer that further reduces the risk of overflow. After chunking, each fragment is submitted as an independent API call with an appended note: "This is part of a larger protocol, extract what you can from this section." The partial results are merged downstream (Section 7).

### 7. Error Handling and Resilience

#### 7.1 Retry with Exponential Backoff and Model Fallback

Network requests are wrapped in a retry loop with three retries and an initial delay of 2,000 ms. The delay doubles on each attempt (2 s, 4 s, 8 s), following standard exponential backoff. Three failure modes are handled distinctly:

- **HTTP 429 (Rate Limit):** The service immediately switches to the next fallback model (Qwen3-235B → GPT-OSS-120B). If no fallback remains, it waits 5 seconds and retries. This automatic failover prevents a single saturated endpoint from blocking an entire batch conversion.
- **HTTP 5xx (Server Error):** Transient server errors trigger the exponential backoff without model switching, under the assumption that the outage is infrastructure-wide rather than model-specific.
- **Network Error / Timeout:** Fetch exceptions (DNS failure, TCP reset, CORS block) are retried with the same backoff curve. All retries are logged to the browser console with timestamps, enabling post-hoc diagnosis.

Client errors (HTTP 4xx except 429) are not retried because they indicate malformed requests, authentication failures, or invalid model IDs that will not resolve with repetition. Instead, the error is surfaced to the user immediately via a toast notification.

#### 7.2 JSON Repair Pipeline

Despite low temperature and explicit formatting instructions, models occasionally emit malformed JSON. A six-stage repair pipeline attempts to salvage the response before discarding it:

1. **Stage 1:** Remove trailing commas before closing brackets or braces.
2. **Stage 2:** Insert missing commas between adjacent objects or arrays.
3. **Stage 3:** Insert missing commas between object properties separated only by newlines.
4. **Stage 4:** Truncate any trailing text after the final closing brace.
5. **Stage 5:** Clean up spacing around closing braces in arrays.
6. **Stage 6:** Add commas before subsequent properties when a closing brace is immediately followed by a quote.

Each repair is applied sequentially, and the result is validated with JSON.parse. If all stages fail, the chunk is skipped and the remaining chunks are still merged. This graceful degradation means that a single malformed section does not invalidate an entire multi-page protocol.

### 8. Output Schema and ISA Generation

The LLM returns a JSON object with two top-level keys: samples and protocols. Samples carry name, organism, and an array of characteristics (category, value, unit, termSource, termAccession). Protocols carry name, description, inputs, parameters, outputs, and dataFiles. This schema was co-designed with the ARCtrl library (v3.0.1) so that the parsed object can be fed directly into ARC JavaScript bindings without intermediate normalisation.

The ISA generation module creates a multi-sheet Excel workbook for each assay or study. Sheet 1 is the sample table, containing Source Name, Organism, and one column per characteristic category. Subsequent sheets are process tables named "process nr. 1", "process nr. 2", and so on. Each process table encodes the ISA-Tab triad: Input → Protocol REF → Output. Parameters appear as additional columns between Protocol REF and Output, using unitised cells when the LLM supplied both a value and a unit. Data files are written as Output [Data] cells with a dataset/ prefix, satisfying the ARC specification that raw data paths be relative to the assay root.

A critical design choice is sequential protocol linking: the outputs of process N are copied into the inputs of process N+1 before table creation. This enforces biological continuity within the ISA representation. If the LLM omits explicit linking, the fallback logic still generates a valid table chain, albeit with generic sample names that can be manually curated later.

#### 9. Hallucination Mitigation

Hallucination — the invention of entities not present in the source text — is the principal risk in automated protocol extraction. Two architectural measures address it.

- **Schema default hardening.** The dataFiles field is defined in the JSON schema as an empty array [] rather than a descriptive string like "array of file names". Early prototypes used the descriptive form and observed 100 % hallucination: every model invented plausible but fictitious filenames (e.g., "results.csv", "FASTQ files"). The empty-array default acts as a null-object pattern; the model only populates the array when filenames are explicitly mentioned in the protocol. After this change, three of four benchmarked models produced zero hallucinated files; the fourth (Gemma-4-31B) persisted with 16 hallucinations, leading to its exclusion from the production allow-list.
- **Explicit array-length rules.** The prompt states that inputs, outputs, and dataFiles must share the same length. This constraint makes spurious entries costly: a model that hallucinates one extra data file must also hallucinate a matching input and output, which is statistically unlikely under low-temperature sampling. The rule therefore acts as a self-consistency check that suppresses low-confidence inventions.

#### 10. Metadata Tracking and Reproducibility

Every conversion that uses the LLM module emits a machine-readable metadata record stored in <assay>/elab2arc-metadata/conversion-{UUID}.json. A parallel latest.json symlink simplifies retrieval. The metadata object contains:

- **Source provenance:** eLabFTW experiment ID, title, author, team, and instance URL.
- **LLM configuration:** Model name, model actually used (after fallback), temperature, max_tokens, and streaming flag.

- **Prompt snapshot:** The complete assembled prompt text, including any user customisations from the Prompt Editor.
- **Chunking telemetry:** Whether chunking was required, the number of chunks, and the estimated token count per chunk.
- **Extraction results:** Number of samples and protocols extracted, plus arrays of errors and warnings.
- **Timing:** Start timestamp, end timestamp, and duration in milliseconds.

Old conversion files are automatically pruned when the directory exceeds ten entries, keeping the ARC lean while preserving a short audit trail. The metadata schema is versioned (currently v1.0.0) and validated on load; missing required fields are flagged with severity levels (error, warning, info). A built-in troubleshooting report generator formats the JSON into Markdown suitable for GitHub issues or email support, dramatically reducing the time required to diagnose user-reported extraction failures.

#### 11. Security and Privacy Considerations

Experimental protocols may contain sensitive information: patient identifiers, proprietary reagent formulations, or pre-publication methodologies. The multi-provider architecture directly addresses these concerns by allowing entirely local inference. When LM Studio or Ollama is selected, protocol text never leaves the user's machine. For remote providers, only the protocol text and the assembled prompt are transmitted; no filesystem paths, git history, or unrelated ARC contents are included in the payload. API keys are stored in browser localStorage, which is sandboxed per origin and never synchronised to cloud backups. The DataPlan proxy adds an institution header so that traffic can be rate-limited and audited at the organisational firewall.

#### 12. Summary of Design Advantages

The LLM function in elab2ARC is engineered for production scientific workflows. The combination of a low-temperature, streaming, chunked extraction pipeline with automatic model fallback and JSON repair yields a 100 % success rate on 350 real-world protocols. The modular prompt system supports reproducible tuning without code changes. The unified provider abstraction ensures that researchers can choose the privacy-cost-latency trade-off appropriate to their institution. Finally, comprehensive metadata tracking closes the reproducibility loop: every ARC generated with LLM assistance carries an auditable record of exactly which model, prompt, and parameters produced it, fulfilling the FAIR principles that underpin the ARC ecosystem.
