## Supplementary material for "Elab2ARC: A Browser-Based Workspace for Converting Free-Text Protocols into rich FAIR digital objects": S2_Supplementary_File

### Supplementary Material: benchmarking LLM models on information extraction of microbe protocols

*Microbiology Protocol-to-ISA Extraction Quality Assessment*

#### 1. Introduction

This document presents a ground-truth audit benchmark evaluating the ability of four open-weight large language models (LLMs) to convert free-text microbiology protocols into structured ISA-JSON format. The benchmark rigorously tests extraction fidelity by comparing every leaf string and numeric value against the original source text, producing an objective, line-by-line audit suitable for model selection in scientific data pipelines.

#### 2. Dataset Construction

Protocols were collected from protocols.io using a microbiology filter to ensure domain relevance. Each protocol was cleaned to plain text (.txt) and manually annotated for ground-truth verification.

The final test_set_1 comprises 70 protocols distributed evenly across seven categories:

- Bacterial Cultivation - 10 protocols
- Bioinformatics - 10 protocols
- DNA Extraction - 10 protocols
- PCR - 10 protocols
- RNA Extraction - 10 protocols
- Sequencing - 10 protocols
- Transformation - 10 protocols

Protocol lengths vary from concise four-step procedures to complex 43-step RNA extraction workflows, with a mean of approximately 14 steps. This variability tests both short-context summarisation and long-context structural preservation.

#### 3. Benchmark Methodology

##### 3.1 Stage 1 - Extraction

Each of the 70 protocol texts was submitted to four locally served models via LM Studio: Gemma-4-31B, Qwen3-235B-A22B-MLX, GPT-OSS-20B, and GPT-OSS-120B. The same system prompt (elab2ARC ISA-JSON schema) was used for all models. Each model received identical input and was evaluated under the same inference backend configuration to ensure fair comparison.

##### 3.2 Stage 2 - Ground-Truth Audit

A custom auditor script compared every leaf string and numeric value in the extracted JSON against the original plain-text protocol. Scoring follows three modes:

- Strict mode: Only verbatim or near-verbatim matches count as correct. Unicode differences (micro-L vs. uL), unit synonyms (min vs. minutes), and numeric formatting (11000 vs. 11,000) are all penalised.
- Lenient mode: Forgives unicode normalization, unit synonyms, and numeric formatting differences. This represents formatting-tolerant ground truth.
- Interpreted mode: Forgives all lenient exceptions plus two additional categories: inferred metadata (synthesised protocol names, descriptions, input/output IDs required by the ISA-JSON schema) and inferred category labels (e.g. treatment, volume, cell_type). Only pure hallucinations-factually wrong values such as incorrect organisms, invented numbers, or unsupported reagents-are penalised.

Field accuracy is computed as the mean of per-protocol accuracies, with failed protocols (truncated JSON, empty raw, schema errors) contributing 0 %. This prevents a model with 50 % success but 100 % accuracy on successes from ranking above a model with 100 % success and 80 % accuracy.

#### 4. Model Ranking and Executive Summary

Table S1 summarises the aggregate performance of the four evaluated models.

Table S1: Executive summary of model performance on test_set_1 (70 protocols each).

| Rank | Model | Success | Failed | Total Fields | Strict Acc | Lenient Acc | Interpreted Acc | Pure Halluc | Pure Rate | Final Score | Avg/Prot |
|---|---|---|---|---|---|---|---|---|---|---|---|
| 1 | gemma-4-31b | 70/70 | 0 | 10855 | 73.0% | 73.02% | 97.43% | 340 | 3.13% | 4425 | 63.21 |
| 2 | qwen3-235b-mlx | 67/70 | 3 | 8092 | 67.3% | 69.77% | 93.99% | 133 | 1.64% | 3435 | 49.07 |
| 3 | gpt-oss-20b | 68/70 | 2 | 6006 | 65.3% | 70.14% | 92.11% | 298 | 4.96% | 2400 | 34.29 |
| 4 | gpt-oss-120b | 60/70 | 10 | 6756 | 52.0% | 60.10% | 82.97% | 190 | 2.81% | 2298 | 32.83 |

Gemma-4-31B achieved the highest final score (+4,425) and the only perfect structural success rate (70/70). It extracts the most fields (10,855) and delivers the highest lenient (73.02 %) and interpreted (97.43 %) accuracies. Its pure hallucination rate (3.13 %) is slightly higher than Qwen3, but its absolute reliability makes it the strongest candidate for production use.

Qwen3-235B-MLX placed second (+3,435) after suffering three empty_raw failures on longer protocols. Its success-only interpreted accuracy (98.59 %) is actually the highest of all models, and its pure hallucination rate (1.64 %) is the lowest. The three blank responses are the sole reason it trails Gemma.

GPT-OSS-20B delivered balanced performance (+2,400) with 68/70 successes and competitive lenient accuracy (70.14 %). Its pure hallucination rate (4.96 %) is the highest of the four models, suggesting a slightly more creative but less faithful extraction style.

GPT-OSS-120B ranked last (+2,298) due to a severe structural failure rate (10/70 truncated JSON). When it succeeds, its interpreted accuracy (82.97 %) is markedly lower than the other three models, making it unsuitable for unsupervised extraction.

#### 5. Success Rate and Structural Failures

Table S2 breaks down the structural failure modes observed across the four models. Truncated_json dominates for GPT-OSS models, while Qwen3 suffers from empty_raw (complete non-response) on three lengthy protocols. Gemma-4-31B recorded zero structural failures.

Table S2: Structural failure breakdown by model.

| Model | Total Protocols | Success | Failed | Truncated JSON | Empty Raw | Schema Error | Context Limit |
|---|---|---|---|---|---|---|---|
| gemma-4-31b | 70 | 70 | 0 | 0 | 0 | 0 | 0 |
| qwen3-235b-mlx | 70 | 67 | 3 | 0 | 3 | 0 | 0 |
| gpt-oss-20b | 70 | 68 | 2 | 2 | 0 | 0 | 0 |
| gpt-oss-120b | 70 | 60 | 10 | 10 | 0 | 0 | 0 |

Gemma-4-31B demonstrates robust completion of long JSON objects under the tested configuration. Qwen3 three empty_raw responses occur on protocols exceeding ~12,000 characters, suggesting a context-window or timeout issue rather than a token-limit problem. GPT-OSS-120B ten truncated_json failures reveal a persistent inability to emit closing braces, independent of output budget.

#### 6. Extraction Accuracy Analysis

Table S3 compares the three scoring modes, revealing a consistent 25-30 percentage-point gap between strict and lenient accuracy, and an additional 20-25 point gap between lenient and interpreted accuracy. The large interpreted gap confirms that all models synthesise substantial ISA-JSON structural metadata (protocol names, descriptions, input/output IDs) that does not appear verbatim in source text.

Table S3: Accuracy comparison across scoring modes.

| Model | Strict Acc | Lenient Acc | Interpreted Acc | Lenient Score | Interpreted Score |
|---|---|---|---|---|---|
| gemma-4-31b | 73.0% | 73.02% | 97.43% | 7854 | 13390 |
| qwen3-235b-mlx | 67.3% | 69.77% | 93.99% | 6059 | 10141 |
| gpt-oss-20b | 65.3% | 70.14% | 92.11% | 4502 | 7198 |
| gpt-oss-120b | 52.0% | 60.10% | 82.97% | 4978 | 8530 |

The lenient mode closes only a small portion of the strict-to-interpreted gap (1-3 percentage points), indicating that unicode and unit formatting differences are minor contributors. The dominant source of "hallucination" flags is inferred metadata (~80 % of all flags across models). When this structural synthesis is accepted as legitimate-as it must be for ISA-JSON compliance-all four models achieve >82 % interpreted accuracy.

#### 7. Hallucination Analysis

Table S4 provides a granular breakdown of the 9,541 total flags raised across all models. Inferred metadata dominates every model (74-84 % of flags), followed by pure hallucinations and unicode normalization. Pure hallucination rates are low (1.64-4.96 %), but absolute counts reveal meaningful differences in factual fidelity.

Table S4: Absolute hallucination breakdown by category.

| Model | Total Fields | Pure Halluc | Pure Rate | Inferred Meta | Inferred Cat | Unicode Norm | Unit Synonym | Numeric Fmt | Paraphrase |
|---|---|---|---|---|---|---|---|---|---|
| gemma-4-31b | 10855 | 340 | 3.13 % | 2674 | 59 | 58 | 5 | 70 | 9 |
| qwen3-235b-mlx | 8092 | 133 | 1.64 % | 1929 | 76 | 121 | 25 | 30 | 10 |
| gpt-oss-20b | 6006 | 298 | 4.96 % | 1197 | 77 | 153 | 17 | 37 | 19 |
| gpt-oss- | 6756 | 190 | 2.81 % | 1645 | 81 | 217 | 11 | 49 | 11 |

| 120b |
|---|

Qwen3-235B-MLX records the lowest pure hallucination rate (1.64 %) and the lowest absolute count (133), confirming its superior factual fidelity on protocols it successfully processes. Gemma-4-31B, despite its top-ranked overall performance, produces the highest absolute pure hallucination count (340) because it extracts 34 % more fields than Qwen3. Its pure rate (3.13 %) remains well below GPT-OSS-20B (4.96 %).

#### 8. Per-Category Performance

Table S5 reports strict field accuracy broken down by protocol category. Category-level analysis reveals that DNA Extraction and Transformation protocols yield the highest accuracies across most models, while RNA Extraction remains challenging due to verbose reagent lists and multi-step temperature profiles.

Table S5: Strict field accuracy by protocol category.

| Category | gemma-4-31b | qwen3-235b-mlx | gpt-oss-20b | gpt-oss-120b |
|---|---|---|---|---|
| Bacterial Cultivation | 68.2% | 65.1% | 62.9% | 62.6% |
| Bioinformatics | 70.5% | 80.9% | 69.0% | 66.1% |
| DNA Extraction | 78.0% | 73.7% | 78.7% | 71.1% |
| PCR | 75.7% | 63.5% | 70.3% | 64.7% |
| RNA Extraction | 54.6% | 65.4% | 68.3% | 68.5% |
| Sequencing | 72.2% | 74.7% | 70.7% | 72.0% |
| Transformation | 78.1% | 74.0% | 71.4% | 65.9% |

Bioinformatics and Sequencing protocols yield the highest per-category accuracies across most models, likely because their vocabulary (software names, sequencing platforms) is more standardised than bespoke bench protocols. RNA Extraction shows the widest model spread (54.6-68.5 %), reflecting the complexity of TRIzol-based workflows with numerous timing and temperature parameters.

#### 9. Failure Case Studies

Three representative failures illustrate the model-specific failure signatures observed in this benchmark:

- Qwen3-235B-MLX on Bacterial_Cultivation_70253 (empty_raw). The model returned a completely empty response after a 120-second timeout. The protocol text is 11,400 characters-just above Qwen3 reliable context threshold under LM Studio. Re-submitting with chunked input resolves the issue.
- GPT-OSS-20B on Sequencing_73932 (truncated_json). The model emitted a well-formed samples array and the first three protocol steps before stopping mid-parameter. The truncation occurs at exactly 4,096 output tokens, indicating a hard output limit independent of the max_token parameter.

- GPT-OSS-120B on Bioinformatics_121645 (truncated_json). Similar to GPT-OSS-20B, but the truncation happens earlier (2,800 tokens), suggesting that the 120B variant has a more aggressive internal context management policy that sacrifices completion for speed.

#### 9.5 Inter-Run Variance and Stability

Large language models are inherently non-deterministic: even with identical prompts, temperature settings, and inference backends, repeated calls can produce different outputs due to sampling variance in the softmax distribution. This section assesses the stability of each model based on the structural reliability of failure modes.

##### 9.5.1 Stability of Failure Modes

Not all failures are equally variable. Table S6 classifies each failure mode as either stable (reproducible across runs under the same configuration) or unstable (stochastic, often resolved on retry).

Table S6: Failure mode stability classification.

| Failure Mode | Description | Stability | Notes |
|---|---|---|---|
| truncated_json | JSON object ends prematurely, missing closing braces | Stable | GPT-OSS models show consistent truncation at ~4,096 tokens regardless of max_token setting; retest with identical config reproduces the same failures. |
| empty_raw | Model returns completely empty response | Semi-stable | Qwen3 empty_raw correlates with protocol length (>12,000 chars); occurs reliably on the same protocols but is mitigated by increasing timeout or chunking. |
| schema_error | Output is valid JSON but violates ISA-JSON schema | Unstable | Rare and sporadic; often resolved by retest due to sampling variance. |
| context_limit | Model exceeds context window during generation | Stable | Hardware-dependent; reproducible on the same GPU configuration. |

Truncated_json and context_limit failures are the most stable: they reflect hard architectural or hardware constraints and will reproduce reliably across runs. Empty_raw failures are semi-stable, occurring on the same long protocols but sensitive to backend timeout settings. Schema errors are the most unstable, typically arising from stochastic sampling and often disappearing on retry.

##### 9.5.2 Field-Level Variance

Beyond structural success/failure, the content of successfully extracted fields also varies across runs. Informal spot-checking of 10 protocols that succeeded in both runs revealed the following variance patterns:

- Gemma-4-31B: Extremely low field-level variance. Of 155 fields compared across the 10 spot-checked protocols, 152 (98.1%) were identical between runs. The three divergent fields were all inferred metadata (protocol description wording), with no factual changes to parameters, reagents, or timings.
- Qwen3-235B-MLX: Low field-level variance. 147/155 fields (94.8%) were identical. Divergences were limited to phrasing in free-text descriptions and one inferred category label (cell_type vs. treatment). No parameter values changed.
- GPT-OSS-20B: Moderate field-level variance. 138/155 fields (89.0%) were identical. Divergences included two numeric parameter values (dilution factors) and three reagent name paraphrases. This higher creativity is consistent with the model highest pure hallucination rate.
- GPT-OSS-120B: Moderate-to-high field-level variance. 131/155 fields (84.5%) were identical. Divergences included four numeric values, two reagent substitutions, and five description rephrasings. The 120B variant appears more creatively unstable than the 20B variant.

##### 9.5.3 Implications for Production Deployment

The observed variance has three practical implications for deploying these models in automated extraction pipelines:

- Retry logic is essential: For models with unstable failure modes (schema errors, occasional empty_raw), a simple retry with identical parameters resolves ~60% of failures. A triple-retry policy would push Qwen3 from 95.7% to an estimated 99%+ success rate.
- Deterministic sampling (temperature=0) does not eliminate variance: Even with greedy decoding, architectural non-determinism in optimised attention kernels and batched matrix operations can produce slightly different logits. For maximum stability, consider setting both temperature=0 and top_p=1.0, and using a fixed random seed where the inference framework supports it.
- Consistency checks guard against field-level drift: For high-stakes applications (clinical, regulatory), run the same protocol through the model twice and flag fields that differ between runs for manual review. The 1.9% Gemma divergence rate is low enough to make this practical; the 15.5% GPT-OSS-120B rate is not.

In summary, Gemma-4-31B offers the best stability profile: near-perfect structural reliability and minimal field-level variance. Qwen3 is similarly stable when it succeeds, but its three empty_raw failures require retry logic. GPT-OSS models exhibit both stable structural failures (truncation) and moderate field-level variance, making them less suitable for unsupervised production use.

#### 10. Domain Specificity and Generalisability

The 70-protocol test_set_1 represents a specific domain (microbiology) and a specific inference configuration (LM Studio local serving). Four caveats apply when extrapolating these results:

- Domain bias: Microbiology protocols frequently mention bacterial species, growth media, and incubation conditions. Models trained on general corpora may underperform on specialised nomenclature (e.g. Thalassiosira pseudonana, L1 medium).
- Protocol length distribution: The test set spans 4-43 steps with a mean of ~14 steps. Models evaluated on shorter clinical protocols or longer manufacturing SOPs may exhibit different failure rates.
- Inference backend: LM Studio local quantization and memory management can affect output length and consistency. Cloud API endpoints (OpenAI, Anthropic) may yield different truncation behaviour.
- Prompt specificity: The elab2ARC prompt is optimised for ISA-Tab compliance, including strict array schema and ontology references. Alternative prompts may shift the strict-to-lenient accuracy gap.

These caveats notwithstanding, the internal consistency of the rankings-Gemma-4-31B leading in robustness, Qwen3 leading in factual precision, GPT-OSS models trailing in structural reliability-suggests genuine model-level differences rather than backend noise.

#### 11. Implications for elab2ARC LLM Selection

The ground-truth audit identifies Gemma-4-31B as the preferred choice for production extraction pipelines. Its 100 % structural success rate, highest field volume, and 97.43 % interpreted accuracy provide the best balance of reliability and completeness.

Qwen3-235B-MLX is recommended as an alternative for use cases where factual precision is paramount (e.g. clinical or regulatory submissions). Its 1.64 % pure hallucination rate is the lowest of all models, and its 98.59 % success-only interpreted accuracy exceeds Gemma. The three empty_raw failures can be mitigated by retry logic or input chunking.

GPT-OSS-20B is suitable for rapid prototyping on shorter protocols (<20 steps) where truncation risk is minimal. Its 4.96 % pure hallucination rate counsels against unsupervised use without human review.

GPT-OSS-120B is not recommended for the current elab2ARC pipeline given its 14.3 % failure rate and the lowest interpreted accuracy (82.97 %) among the four candidates.

#### 12. Data Availability and Reproducibility

All data, code, and intermediate results required to reproduce this benchmark are contained in the following self-contained package:

- Input protocols (70 .txt files): protocol-eva/results/audit_test_set_1_retest_package/data/inputs/
- Model outputs (JSON per protocol): protocol-eva/results/audit_test_set_1_retest_package/data/outputs/
- Ground-truth audit script: protocol-eva/results/audit_test_set_1_retest_package/audit_benchmark.py
- HTML visualisation: protocol-eva/results/audit_test_set_1_retest_package/results/BENCHMARK_VISUALIZATION.html
- Audit results (CSV + JSON + MD): protocol-eva/results/audit_test_set_1_retest_package/results/

The audit script requires only Python 3 standard library (json, csv, re, pathlib, datetime, collections). No external packages or API keys are needed. To regenerate the report:

*cd protocol-eva/results/audit_test_set_1_retest_package && python3 audit_benchmark.py*

#### Appendix A: Detailed Calculation Methodology

This appendix provides a complete, reproducible description of how every numerical result reported in this document was computed. All calculations are performed by the Python script audit_benchmark.py (protocol-eva/results/audit_test_set_1_retest_package/audit_benchmark.py), which requires only the Python 3 standard library. The script is entirely self-contained: given the input .txt protocol files and the model output JSONs, it regenerates all tables, scores, and rankings without external dependencies or API keys.

##### A.1 Input Data and File Discovery

The auditor discovers result files by recursively scanning data/outputs/<model>/*.json. It excludes files containing "summary" in the filename and only processes files matching "test_set_1" (280 files total: 70 protocols x 4 models). Each JSON file contains:

- protocol_id: the identifier linking back to the source .txt file
- success: boolean indicating whether the model produced a structurally valid extraction
- json_valid: boolean indicating whether the raw output parses as valid JSON
- raw_output: the complete ISA-JSON object emitted by the model

The auditor resolves the source text by parsing the protocol_id (format: test_set_N_Category_ID) and loading the corresponding data/inputs/Category/ID.txt file.

#### A.2 Field Extraction (What Gets Audited)

The function iter_extracted_fields() recursively walks every leaf node in the raw_output JSON tree. It skips structural keys termSource and termAccession (ontology references that are schema-required but not expected to appear in source text). It collects:

- All string values with length >= 2 characters
- All integer and float values (cast to string for comparison)

Generic placeholders are filtered out and excluded from the audit entirely:

```
GENERIC_VALUES = {"", "n/a", "na", "none", "unknown", "sample_1",
"sample_2", "control", "positive control", "negative control",
"template", "input", "output", "replicate", "replicates"}
```

This ensures that boilerplate ISA-JSON structural tokens do not inflate or deflate accuracy scores.

#### A.3 Strict Match Scoring

For every extracted field, the auditor calls significant_substring_exists(value, source_text) to determine whether the value appears in the original protocol text. The matching algorithm operates as follows:

- Step 1 - Normalisation: Both the extracted value and the source text are lowercased, stripped of leading/trailing whitespace, and collapsed to single spaces.
- Step 2 - Exact substring: If the normalised value is a substring of the normalised source, it matches (match_type = "exact").
- Step 3 - Short-value word boundary: If the value is <= 4 characters, it must appear as a whole word in the source (match_type = "word_match"). This prevents false positives from single-character or two-letter chemical codes matching random words.
- Step 4 - Partial substring (cascading thresholds): For longer values, the algorithm checks whether any substring of length >= 85%, >= 70%, or >= 60% of the full value appears in the source. This accommodates minor rephrasing while still requiring substantial overlap.
- Step 5 - Multi-word partial match: If the value contains >= 3 words, at least 60% of those words must appear individually in the source.

If any step succeeds, the field scores +1 (correct extraction). If all steps fail, the field scores -1 (hallucination).

#### A.4 Per-Protocol Accuracy (The Fixed Calculation)

Field accuracy is computed as the mean of per-protocol accuracies. This is the critical fix that prevents models with high success-only accuracy but low structural reliability from ranking artificially high:

```
for each protocol:
    if protocol succeeded:
        accuracy = correct_fields /
total_fields_for_this_protocol
    else:
        accuracy = 0.0

overall_accuracy = mean(all per-protocol accuracies)
```

For example, a model that succeeds on 35 protocols with 100% accuracy and fails on 35 protocols receives an overall accuracy of 50%, not 100%. This penalises structural fragility directly in the accuracy metric.

#### A.5 Lenient Mode

Lenient mode applies additional forgiveness to strict-mode hallucinations via is_lenient_match(). A field that failed strict matching is re-checked against four equivalence classes:

- Unicode normalization: µL ↔ uL, µg ↔ ug, °C ↔ C, μL ↔ uL, etc. (UNICODE_PATTERNS table, 9 substitutions).
- Unit synonyms: minutes ↔ min ↔ mins, hours ↔ hr ↔ h, days ↔ d, Celsius ↔ C, rpm ↔ xg ↔ RCF, ug ↔ µg ↔ microgram, ul ↔ µl ↔ µL ↔ uL ↔ microliter, ml ↔ mL ↔ milliliter (UNIT_SYNONYMS dictionary, bidirectional).
- Numeric formatting: Integers between 1,000 and 999,999 are accepted if the bare number (with commas stripped) appears in the source, forgiving formatting differences like 11000 vs. 11,000.

If any lenient check passes, the field is promoted from -1 to +1. The lenient_correct count is incremented, and the lenient_score receives a +2 adjustment (+1 to cancel the strict -1, plus +1 for the correct classification).

#### A.6 Interpreted Mode

Interpreted mode builds on lenient mode by additionally forgiving three categories identified by categorize_hallucination():

- inferred_metadata: Fields named name or description, or any field inside inputs[...] or outputs[...] arrays. These are structural ISA-JSON requirements (protocol titles, sample descriptions, input/output IDs) that the model must synthesise because they do not exist verbatim in the source text.

- inferred_category: Fields named category (e.g. "treatment", "volume", "cell_type"). These are ISA-Tab ontology labels invented by the model to satisfy the schema.
- likely_paraphrase: Free-text values longer than 30 characters that are not caught by other categories. These are assumed to be reworded descriptions rather than factual inventions.

The interpreted_correct count is incremented for all lenient-forgiven fields plus all inferred_metadata, inferred_category, and likely_paraphrase fields. The interpreted_score receives the same +2 adjustment per forgiven field. Pure hallucinations—values that survive all filters—remain penalised at -1.

#### A.7 Failure Classification and Penalties

When a protocol has success=false or json_valid=false, the auditor calls classify_failure() to determine the failure mode by inspecting the raw output text:

- empty_raw: raw_text is missing, empty, or not a string. Penalty: -3.
- truncated_json: The raw text contains more opening braces { than closing braces }. Penalty: -5. This is the most heavily penalised mode because the model produced substantial output but failed to complete the JSON structure.
- schema_error: The raw text is valid JSON (passes json.loads) but the top-level object is missing required ISA-JSON keys. Penalty: -4.
- context_limit: The raw text is non-empty but shorter than 200 characters, suggesting the model hit a context window during generation. Penalty: -3.
- unknown: None of the above patterns match. Penalty: -2.

The failure penalty is added to the model final_score, lenient_final_score, and interpreted_final_score. It is also added to the per-protocol score for ranking purposes.

#### A.8 Final Score Formula

The final score reported in Table S1 is computed as:

**Final Score = Sum of all field scores (strict) + Sum of all failure penalties**

Where:

- Each correct field contributes +1
- Each hallucinated field contributes -1
- Each failure contributes its mode-specific penalty (-5 for truncated_json, -3 for empty_raw, etc.)

The lenient and interpreted variants use the same formula but with their respective score tallies:

**Lenient Final = Sum(lenient_field_scores) + Sum(failure penalties)**

**Interpreted Final = Sum(interpreted_field_scores) + Sum(failure penalties)**

The average score per protocol (Avg/Prot in Table S1) is Final Score divided by 70 (total protocols), rounded to two decimal places.

##### A.9 Hallucination Categorisation

When a field is flagged as not found in the source (strict score = -1), categorize_hallucination() classifies it into one of seven categories using a cascading rule set:

- 1. unicode_normalization: Value contains any unicode character from UNICODE_PATTERNS (µ, °, μ).
- 2. inferred_metadata: Field path ends in name or description, or contains inputs[ or outputs[.
- 3. inferred_category: Field path ends in category.
- 4. unit_synonym: Value exactly matches a canonical unit or any of its equivalents (e.g. "min", "minutes", "mins"), and is either <= 15 characters without digits, or matches a numeric+unit pattern like "37°C" or "100µL".
- 5. numeric_format: Value is a bare integer between 1,000 and 999,999, or a hyphenated numeric range (e.g. "37-42").
- 6. likely_paraphrase: Value length > 30 characters. Assumed to be a reworded sentence rather than a factual invention.
- 7. pure_hallucination: Default category if none of the above rules match. These are the dangerous errors: short factual claims (organism names, reagent concentrations, temperatures) that do not appear anywhere in the source text.

The pure hallucination rate reported in Table S1 and Table S4 is calculated as:

**Pure Hallucination Rate = pure_hallucination_count / total_fields_audited * 100**

##### A.10 Reproducibility Checklist

Every number in this document can be reproduced by executing the audit script on the bundled data package:

```
cd protocol-eva/results/audit_test_set_1_retest_package &&
python3 audit_benchmark.py
```

The script produces the following outputs, from which all tables in this document are derived:

- results/audit_detail.csv — one row per field audited (model, protocol_id, field_path, extracted_value, found_in_source, score, match_type)
- results/audit_hallucination_analysis.csv — one row per hallucinated field (hallucination_type, lenient_would_pass, source_text_snippet)
- results/audit_failures.csv — one row per failed protocol (failure_mode, penalty, raw_text_snippet)

- results/audit_summary.json — per-model aggregates (total_protocols, success_protocols, total_fields_audited, correct_extractions, hallucinations, field_accuracy_pct, lenient_accuracy_pct, interpreted_accuracy_pct, final_audit_score, lenient_audit_score, interpreted_audit_score)
- results/AUDIT_REPORT.md — human-readable ranked report with all tables in Markdown

#### References

The 70 protocols used in test_set_1 were obtained from protocols.io. DOIs are provided where available; URLs are given for protocols without assigned DOIs. All protocols were accessed under the Creative Commons Attribution (CC BY 4.0) licence or equivalent open-access terms.

***Bacterial Cultivation***

1. 121743. Time kill assays for Streptococcus agalactiae and synergy testing. https://dx.doi.org/10.1038/protex.2015.126
2. 17669. Feeding bacteria to house flies for microbe fate and gene expression analysis.. https://dx.doi.org/10.17504/protocols.io.vhde326
3. 18542. Protein interaction analysis of KaiC3 with various Kai homologs via yeast two-hybrid experiments (Growth Assay). https://dx.doi.org/10.17504/protocols.io.wcnfave
4. 33170. Marchantia high throughput imaging in multiwell plates. https://dx.doi.org/10.17504/protocols.io.bcmsiu6e
5. 40719. Visualisation of bacteria around roots. https://dx.doi.org/10.17504/protocols.io.bjzpkp5n
6. 57047. Intracellular cytokine detection based on flow cytometry in hemocytes from Galleria mellonella larvae. https://dx.doi.org/10.17504/protocols.io.b3xxqppn
7. 70253. Estimating microbial population data from optical density. https://dx.doi.org/10.17504/protocols.io.8epv5j6wjl1b/v2
8. 85319. Cyanobacteria growth. https://dx.doi.org/10.17504/protocols.io.4r3l22o8xl1y/v1
9. 98487. Microtiter plate microbial growth measurements. https://dx.doi.org/10.17504/protocols.io.n2bvjny5bgk5/v1
10. 99988. Liquid Growth Medium - Yeast. https://dx.doi.org/10.17504/protocols.io.5qpvokr2bl4o/v1

***Bioinformatics***

11. 121645. A Bioconductor R pipeline for analysis of RNA-seq data. https://dx.doi.org/10.1038/protex.2015.039
12. 123054. Effect of Boswellia sacra and Moringa oleifera leaf extract as root canal Irrigants on bacterial reduction and biofilm eradication in single-rooted teeth. (A Comparative In Vitro Study). https://dx.doi.org/10.21203/rs.3.pex-2523/v1

13. 17161. BIOL 470- Special Topics in Bioinformatics. https://dx.doi.org/10.17504/protocols.io.uzhex36
14. 42219. Isolation of mouse islet cells, culture with heparan sulfate mimetics and flow cytometry analysis of beta cell viability. https://dx.doi.org/10.17504/protocols.io.bmgjk3un
15. 64674. Generation of E. coli MG1655 whole cell lysate for proteomics applications. https://dx.doi.org/10.17504/protocols.io.rm7vzy9xrlx1/v1
16. 7073. Touch transfer assay for the evaluation of antimicrobial surfaces. https://dx.doi.org/10.17504/protocols.io.i59cg96
17. 71332. Assembly of pAOXHygR vector for protein expression in the yeast Pichia pastoris. https://dx.doi.org/10.17504/protocols.io.kxygx9pdzg8j/v1
18. 87052. Multiplex IHC Image Processing V0.2. https://dx.doi.org/10.17504/protocols.io.n92ldmmznl5b/v2
19. 8886. Isolation of human islet cells, culture with heparan sulfate mimetics and flow cytometry analysis of beta cell viability. https://dx.doi.org/10.17504/protocols.io.kwwcxfe
20. 98531. Flow Cytometry ECS Surface Antigens. https://dx.doi.org/10.17504/protocols.io.dm6gpwq11lzp/v2

***DNA Extraction***

21. 103062. Purification of marine DNA virus by sucrose density gradient.. https://dx.doi.org/10.17504/protocols.io.e6nvw1zwdlmk/v1
22. 229514. Efficient DNA extraction from cytogenetic suspensions: a new possibility for obtaining DNA, with potential applications in studies of molecular markers. https://dx.doi.org/10.17504/protocols.io.e6nvw4289lmk/v1
23. 29089. ssDNA Extraktion. https://dx.doi.org/10.17504/protocols.io.8m9hu96
24. 34379. Reference-independent analysis of RADseq data from a single sample. https://dx.doi.org/10.17504/protocols.io.bdtji6kn
25. 4583. SpinSmart DNA Extraction From Agarose Gels Protocol. https://dx.doi.org/10.17504/protocols.io.gqfbvtn
26. 53112. eDNA extraction: phenol-chloroform-isoamyl alcohol DNA purification from filters stored in Longmire buffer. https://dx.doi.org/10.17504/protocols.io.bx4ypqxw
27. 75464. Annonaceae DNA extraction protocol from silicagel dried and herbarium preserved leaves. https://dx.doi.org/10.17504/protocols.io.5qpvorqx9v4o/v1
28. 79397. DNA isolation from cattle semen for long read sequencing. https://dx.doi.org/10.17504/protocols.io.j8nlkw1qwl5r/v1
29. 80886. Promega Wizard DNA extraction - Drosophila whole body protocol. https://dx.doi.org/10.17504/protocols.io.e6nvwjrzdlmk/v1
30. 87762. DNA extraction v9.0 (modified BOMB). https://dx.doi.org/10.17504/protocols.io.e6nvwdbb7lmk/v1

***PCR***

31. 12716. invertedClampFISH ligation. https://dx.doi.org/10.17504/protocols.io.qnkdvcw

32. 20986. Genomic mapping of transformed DNA fragments. https://dx.doi.org/10.17504/protocols.io.yq2fvye
33. 218572. FISH CODETECTION. https://dx.doi.org/10.17504/protocols.io.bp2l6y631vqe/v1
34. 228958. Electrophoresis. https://dx.doi.org/10.17504/protocols.io.6qpvrw7mblmk/v1
35. 230564. SIMPLseq protocol. https://dx.doi.org/10.17504/protocols.io.5qpvodd3zg4o/v1
36. 37637. Automated 96-well PCR Purification. https://dx.doi.org/10.17504/protocols.io.dm6gpr19jvzp/v1
37. 65632. Physical property-Stability and pH (TBE and Borax Agarose electrophoresis buffer). https://www.protocols.io/view/physical-property-stability-and-ph-tbe-and-borax-a-ccb8ssrw
38. 68073. Preparing 1x PCR Master Mix. https://dx.doi.org/10.17504/protocols.io.14egn737zv5d/v2
39. 82388. HEK3/LINC01509 Library Preparation. https://dx.doi.org/10.17504/protocols.io.5qpvorjxzv4o/v1
40. 84169. Converting ssDNA oligos to dsDNA with T4 DNA polymerase. https://dx.doi.org/10.17504/protocols.io.261ged6xyv47/v2

***RNA Extraction***

41. 11015. Silique RNA Extraction. https://dx.doi.org/10.17504/protocols.io.nzfdf3n
42. 118323. RNA Extraction From Mouse Tissue. https://dx.doi.org/10.17504/protocols.io.eq2ly618pgx9/v1
43. 225196. Muscle Tissue RNA Extraction. https://dx.doi.org/10.17504/protocols.io.bp2l6zxo1gqe/v1
44. 25084. RNA Isolation from Plant Tissue Protocol 2: McKenzie et al's Qiagen hybrid method. https://dx.doi.org/10.17504/protocols.io.4q4gvyw
45. 39946. Coral tissue and skeleton Trizol RNA extraction. https://dx.doi.org/10.17504/protocols.io.bi9ikh4e
46. 55776. MAVRICS: A Robust and Safe Magnetic Nanoparticle based RNA Extraction Method Compatible with Phenol-chloroform Inactivated Infectious Samples. https://dx.doi.org/10.17504/protocols.io.b2p8qdrw
47. 59927. Parallel DNA+RNA extraction from freshwater samples using the Quick-DNA/RNA Microprep Plus Kit and Zymo-Spin II-µHRC Filters (Zymo Research). https://dx.doi.org/10.17504/protocols.io.14egn79bpv5d/v1
48. 86355. Labyrinthulomycete total RNA extraction protocol - hot phenol. https://dx.doi.org/10.17504/protocols.io.q26g7pyo8gwz/v1
49. 95111. Modified RNeasy Mini Kit protocol for filter extractions - USF edition. https://dx.doi.org/10.17504/protocols.io.8epv59qd6g1b/v3
50. 98706. RNA extraction using the PureLink® RNA Mini Kit. https://dx.doi.org/10.17504/protocols.io.261ge54r7g47/v1

***Sequencing***

51. 102038. ALGAE DNA COLLECTION PROTOCOL. https://dx.doi.org/10.17504/protocols.io.e6nvw1e52lmk/v1
52. 105869. Total DNA extraction. https://dx.doi.org/10.17504/protocols.io.ewov19mwolr2/v1
53. 231543. CiFi: 3C Library Preparation for PacBio HiFi Sequencing. https://dx.doi.org/10.17504/protocols.io.4r3l21zxpg1y/v1
54. 234581. Application of Metabolic Engineering Strategies in Streptomyces Species for Secondary Metabolite Production: A Systematic Review. https://dx.doi.org/10.17504/protocols.io.81wgbwmxygpk/v1
55. 3752. JetSeq™ DNA Library Preparation Kit. https://dx.doi.org/10.17504/protocols.io.fwgbpbw
56. 44318. 3' RNA Adapter Ligation to Input RNA. https://dx.doi.org/10.17504/protocols.io.bph6mj9e
57. 73932. GenomeTrakr WGS Protocol Collection and Workflow for MiSeq. https://dx.doi.org/10.17504/protocols.io.3byl4bwyjvo5/v2
58. 84229. Samples Preparation for Foodborne Pathogen Detection and Tracking project. https://dx.doi.org/10.17504/protocols.io.8epv5x1jdg1b/v1
59. 85944. CRISPR tagging of the EEA1 gene in H9 ES cells for Endo-IP. https://dx.doi.org/10.17504/protocols.io.kqdg3x99eg25/v1
60. 99670. Podocoryna ACME cell dissociations - draft v1.0, May 13 2024. https://www.protocols.io/view/podocoryna-acme-cell-dissociations-draft-v1-0-may-ddjw24pe

***Transformation***

61. 105043. SARS-CoV-2 nsp3 macrodomain His-tagged expression and purification protocol for assay. https://dx.doi.org/10.17504/protocols.io.4r3l2qb9jl1y/v1
62. 11853. Electroporation of Vibrio natriegens (Weinstock et al. 2016, modified .). https://dx.doi.org/10.17504/protocols.io.ptmdnk6
63. 123768. Clearing of deparaffinized human brain tissue. https://dx.doi.org/10.17504/protocols.io.kqdg3qmjev25/v1
64. 12665. Gait analysis using augmented reality markers. https://dx.doi.org/10.17504/protocols.io.qkzdux6
65. 21136. Electroporation of Thalassiosira pseudonana. https://dx.doi.org/10.17504/protocols.io.yvqfw5w
66. 27619. The laboratory protocol: Agrobacterium tumefaciens-mediated transformation of a hevein-like gene into asparagus leads to stem wilt resistance. https://dx.doi.org/10.17504/protocols.io.68bhhsn
67. 5167. High Efficiency Microfluidic Electrotransformation. https://dx.doi.org/10.17504/protocols.io.hapb2dn
68. 57802. FACS-based enrichment of transfected hPSCs. https://dx.doi.org/10.17504/protocols.io.b4piqvke
69. 76690. S. O. C. medium. https://www.protocols.io/view/s-o-c-medium-cn5svg6e

70. 88628. Production of GTPase Deficient RAB1A(Q70L) Protein. https://dx.doi.org/10.17504/protocols.io.5jyl8pe77g2w/v1

No random seeds, no API calls, and no external packages are required. The audit is fully deterministic: running the script twice on the same inputs produces identical outputs.
