## Supplementary material for "Elab2ARC: A Browser-Based Workspace for Converting Free-Text Protocols into rich FAIR digital objects": S4_Supplementary_File

### Experiment 1: Bacterial Cultivation

**Date:** 2025-07-02
**Tags:** cultivation bacteria
**Category:** Assay
**Status:** Success
**Created by:** elab elab

#### Bacterial cultivation of *E. coli* Strain K-12 (n=10)

#### Goal:

To cultivate *Escherichia coli* K-12 under controlled conditions and monitor growth for subsequent DNA extraction.

#### Procedure:

**Strains:** *E. coli* K-12
**Medium:** LB broth, 10 mL per culture
**Conditions:** 37 °C, 180 rpm, 16 h

**Inoculation:** 1:100 dilution from overnight culture (OD600 = 2.5)

**Sampling:** OD600 measurements taken every hour for 8 hours.

**OD600 measurements (after overnight incubation):**

| Time (Hours) | Sample 1 (OD600) | Sample 2 (OD600) | Sample 3 (OD600) | Sample 4 (OD600) | Sample 5 (OD600) | Sample 6 (OD600) | Sample 7 (OD600) | Sample 8 (OD600) | Sample 9 (OD600) | Sample 10 (OD600) |
|---|---|---|---|---|---|---|---|---|---|---|
| 0 | 0,051 | 0,049 | 0,053 | 0,050 | 0,052 | 0,051 | 0,048 | 0,050 | 0,053 | 0,049 |
| 1 | 0,098 | 0,095 | 0,102 | 0,097 | 0,100 | 0,099 | 0,094 | 0,096 | 0,101 | 0,096 |
| 2 | 0,210 | 0,205 | 0,215 | 0,208 | 0,212 | 0,211 | 0,203 | 0,207 | 0,214 | 0,206 |
| 3 | 0,450 | 0,442 | 0,458 | 0,445 | 0,453 | 0,451 | 0,439 | 0,443 | 0,456 | 0,440 |
| 4 | 0,890 | 0,875 | 0,905 | 0,880 | 0,895 | 0,892 | 0,870 | 0,878 | 0,900 | 0,872 |
| 5 | 1.520 | 1.490 | 1.550 | 1.500 | 1.530 | 1.525 | 1.480 | 1.495 | 1.545 | 1.485 |
| 6 | 1.950 | 1.920 | 1.980 | 1.930 | 1.960 | 1.955 | 1.910 | 1.925 | 1.975 | 1.915 |
| 7 | 2.100 | 2.070 | 2.130 | 2.080 | 2.110 | 2.105 | 2.060 | 2.075 | 2.125 | 2.065 |
| 8 | 2.150 | 2.120 | 2.180 | 2.130 | 2.160 | 2.155 | 2.110 | 2.125 | 2.175 | 2.115 |

**Notes:** Cultures harvested at 6 hours (5,000 g, 10 min, 4 °C) for optimal cell density and viability for DNA extraction.

Unique eLabID: 20250702-db58d72d05b2f043f64e9832453bf63c25d66d34
Link: https://elab.dataplan.top/experiments.php?mode=view&id=40
