## Supplementary material for "Elab2ARC: A Browser-Based Workspace for Converting Free-Text Protocols into rich FAIR digital objects": S5_Supplementary_File

### Experiment 2: DNA Extraction

**Date:** 2025-07-03
**Tags:** DNA extraction bacteria
**Category:** Assay
**Status:** Success
**Created by:** elab elab

---

### Genomic DNA extraction from *E. coli* K-12

### Goal:

Extract DNA of the strains cultivated in Experiment 1: Bacterial Cultivation for further analysis

### Procedure:

Use Methode - NucleoSpin Microbial DNA Kit (Macherey & Nagel) for DNA extraction.

Manufacturer's instructions, with lysozyme pre-treatment (20 mg/mL, 30 min, 37 °C)

- MN Bead Tubes Type B

- Disruption time 4 min

### Results:

**Elution volume:** 100 µL BE buffer

##### DNA concentration (Qubit dsDNA HS Assay):

| SAMPLE ID | DNA CONC. [NG/ML] | A260/280 | A260/230 |
|---|---|---|---|
| EC-01 | 95.5 | 1.81 | 2.10 |
| EC-02 | 87.2 | 1.78 | 2.05 |
| EC-03 | 89.7 | 1.80 | 2.10 |
| EC-04 | 75.7 | 1.79 | 2.08 |
| EC-05 | 90.5 | 1.80 | 2.09 |
| EC-06 | 89.8 | 1.78 | 2.08 |
| EC-07 | 88.1 | 1.81 | 2.10 |
| EC-08 | 97.7 | 1.81 | 2.06 |
| EC-09 | 96.4 | 1.79 | 2.07 |
| EC-10 | 86.6 | 1.78 | 2.10 |

**Storage:** -20 °C until library prep
**Notes:** DNA was of sufficient purity for downstream library preparation.

Quality check:

Gel type: 1% Agarose, TAE, Ethidium Bromide

Ladder: Generuler 1kb Plus

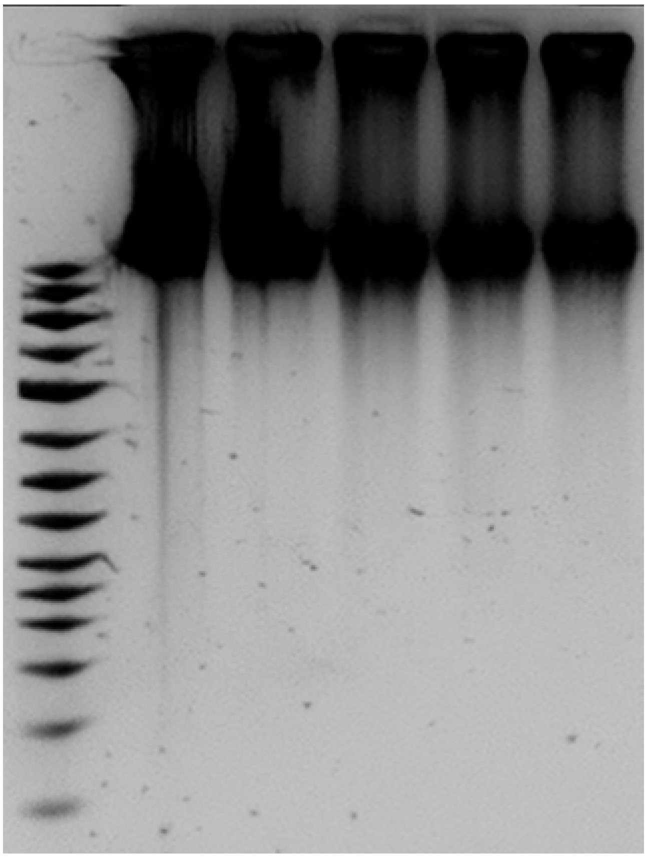

High molecular weight DNA observed for all samples, no signs of shearing or degradation.

#### Linked experiment

Assay - Experiment 1: Bacterial Cultivation

#### Linked resource

Methode - NucleoSpin Microbial DNA Kit (Macherey & Nagel)

### Attached file

2025-07-02_DNAExtractionpicture.png
sha256: c9cbac042324699ea37748184216d7ba8291b6062771ad7e01b17211e1522ffb

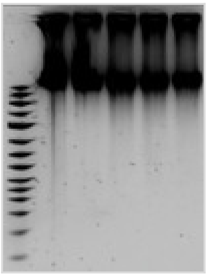

Unique eLabID: 20250702-c2c603c53dc4d4b3aa98b9eb41c50b5953503ea5
Link: https://elab.dataplan.top/experiments.php?mode=view&id=41
