## Supplementary material for "Elab2ARC: A Browser-Based Workspace for Converting Free-Text Protocols into rich FAIR digital objects": S6_Supplementary_File

### Experiment 3: Library Preparation - FFPE Repair, A-tailing, Adapter Ligation

**Date:** 2025-07-04
**Tags:** bacteria library preparation
**Category:** Assay
**Status:** Success
**Created by:** elab elab

#### Goal:

#### Library preparation using NEBNext FFPE DNA Repair Mix

#### Procedure:

**Steps performed:**

- FFPE DNA repair Methode - NEBNext FFPE DNA Repair Mix
- End-repair and A-tailing
- Ligation of Illumina-compatible adapters (NEBNext Multiplex Oligos)
  **Input DNA:** 200 ng per sample ( Assay - Experiment 2: DNA Extraction)
  **Clean-up:** AMPure XP beads (1.8x)
  **QC:** Bioanalyzer traces (average library size ~420 bp)
  **Output:** Indexed libraries for sequencing
  **Storage:** -20 °C
  **Notes:** No signs of adapter dimers; libraries passed QC for sequencing.

#### Results:

**DNA concentration (Qubit dsDNA HS Assay):**

| SAMPLE ID | DNA CONC. [NG/ML] | A260/280 | A260/230 |
|---|---|---|---|
| EC-01 | 50.5 | 1.80 | 2.11 |
| EC-02 | 45.2 | 1.77 | 2.08 |
| EC-03 | 52.5 | 1.81 | 2.06 |
| EC-04 | 44.7 | 1.78 | 2.09 |
| EC-05 | 48.5 | 1.81 | 2.11 |
| EC-06 | 42.8 | 1.78 | 2.10 |
| EC-07 | 35.1 | 1.81 | 2.09 |
| EC-08 | 54.5 | 1.80 | 2.10 |
| EC-09 | 35.2 | 1.77 | 2.07 |
| EC-10 | 45.4 | 1.75 | 2.10 |

#### Linked experiment

Assay - Experiment 2: DNA Extraction

#### Linked resource

Methode - NEBNext FFPE DNA Repair Mix

#### Attached file

Primer_sequences.txt
sha256: 2d3feccd84046b5f4bf4d4af9587413bf7084d36ac1aa3fdb16998751a126a6a

Unique eLabID: 20250702-a5a73030192820062a83814c3026497ecd79241e
Link: https://elab.dataplan.top/experiments.php?mode=view&id=42
