## Supplementary material for "Elab2ARC: A Browser-Based Workspace for Converting Free-Text Protocols into rich FAIR digital objects": S7_Supplementary_File

### Experiment 4: Sequencing Run

**Date:** 2025-07-05
**Tags:** bacteria sequencing
**Category:** Assay
**Status:** Success
**Created by:** elab elab

### Illumina NovaSeq 6000 sequencing

### Goal:

Sequencing of samples from Assay - Experiment 3: Library Preparation - FFPE Repair, A-tailing, Adapter Ligation

### Methods:

- **Sequencing Platform:** Illumina NovaSeq 6000
- **Flow Cell:** SP Flow Cell (Lot No. NV-FC-20250622A)
- **Chemistry:** NovaSeq 6000 S1 Reagent Kit v1.5 (300 cycles)
- **Loading Concentration:** 200 pM
- **Read Length:** 2x150 bp paired-end reads

.

### Results:

- **Yield:** Approximately 10-15 Gb per sample.
- **Q30 Score:** >90% across all samples.
- **Data Location:** FASTQ files are stored on the institutional `smb://institutionserver/microbial_genomics_data/raw_reads/`

#### Linked experiment

Assay - Experiment 3: Library Preparation - FFPE Repair, A-tailing, Adapter Ligation

Unique eLabID: 20250702-0efc561047555f209be2d8d5b40949cbb075318a
Link: https://elab.dataplan.top/experiments.php?mode=view&id=43
