## Supplementary material for "Elab2ARC: A Browser-Based Workspace for Converting Free-Text Protocols into rich FAIR digital objects": S8_Supplementary_File

### Experiment 5: Bioinformatic Analysis

**Date:** 2025-07-06
**Tags:** bacteria sequencing analysis
**Category:** Assay
**Status:** Success
**Created by:** elab elab

#### Goal:

Preprocessing and assembly of *E.coli* genomes

Used data from sequencing run Assay - Experiment 4: Sequencing Run

#### Procedure:

**Methods:**

- **Software:**
  - **Quality Control:** FastQC (v0.11.9), MultiQC (v1.9)
  - **Trimming:** Trimmomatic (v0.39)
  - **Assembly:** SPAdes (v3.15.3)
  - **Annotation:** Prokka (v1.14.6), NCBI RefSeq
- **Bioinformatics Pipeline:** Custom Snakemake workflow (Version 1.2)
- **Reference Database:** NCBI RefSeq bacterial genomes (downloaded 2025-06-01)

**Results:**

- **QC Reports:** All samples passed quality thresholds (high quality reads, minimal adapter contamination).
- **Assembly Statistics:**
  - Average Contig N50: 1.5 Mbp
  - Total Contig Length: ~4.8-5.2 Mbp per sample (consistent with *E. coli* genome size)
  - Number of Contigs: 50-70 per sample
- **Annotation:**
  - Number of Predicted Genes: ~4500-5000 per sample
  - Identified Species: *Escherichia coli* K-12 confirmed for all samples.

#### Results:

**Data Access Link (External):**
`smb://institutionserver/microbial_genomics_data/analysis/20250625_MGL_Analysis_Run1` (Contains assembled genomes, annotation files, QC reports)

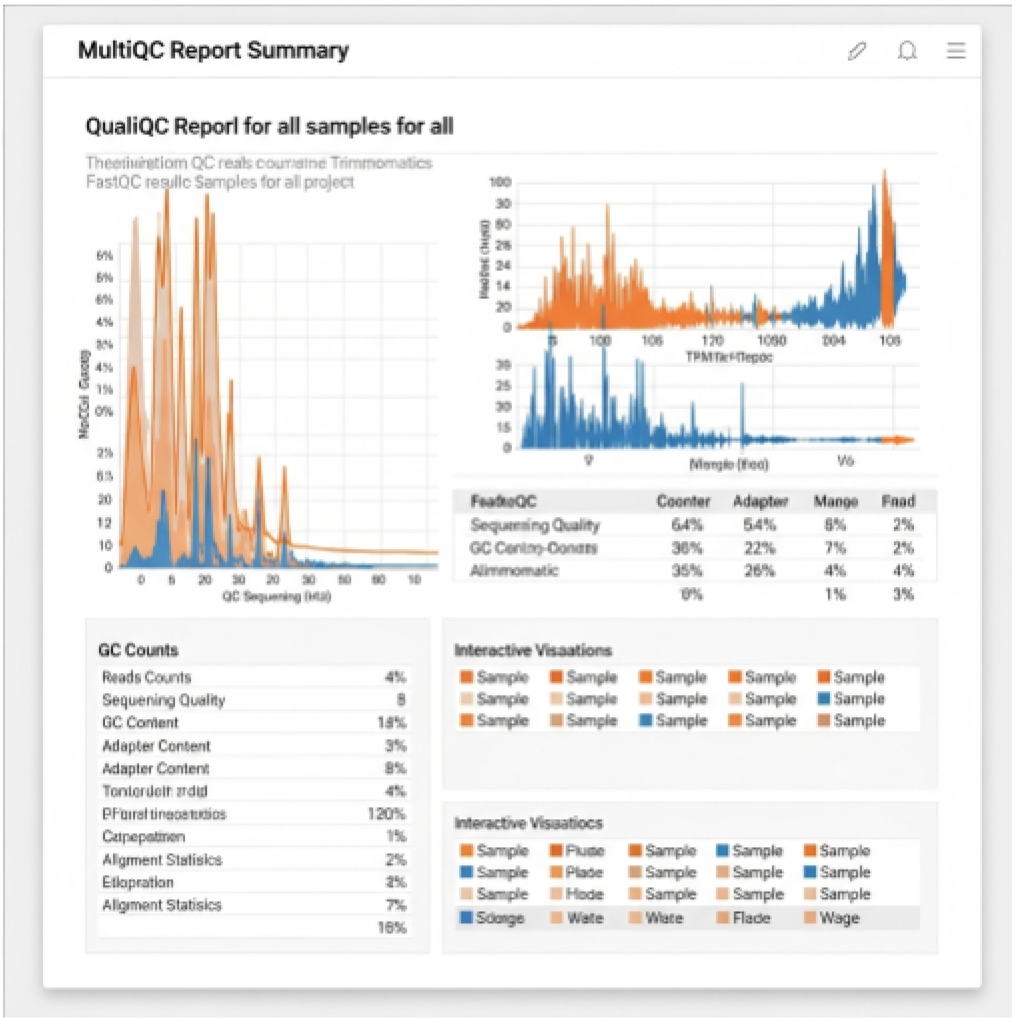


#### Linked experiment

Assay - Experiment 4: Sequencing Run

#### Attached file

Screenshot-2025-06-18-124708.png
sha256: ac52f639d598f84662f5ed556a28554fb24d75f01f222e1d222e12ebe1b326bf

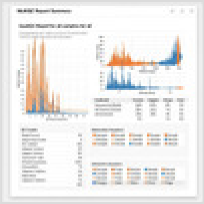

Unique eLabID: 20250702-02e95e6d88ac63d4d27a43f12881005992bf7aa3
Link: https://elab.dataplan.top/experiments.php?mode=view&id=44
